# Scalable ultra-high-throughput single-cell chromatin and RNA sequencing reveals gene regulatory dynamics linking macrophage polarization to autoimmune disease

**DOI:** 10.1101/2023.12.26.573253

**Authors:** Sara Lobato-Moreno, Umut Yildiz, Annique Claringbould, Nila H. Servaas, Evi P. Vlachou, Christian Arnold, Hanke Gwendolyn Bauersachs, Víctor Campos-Fornés, Karin D. Prummel, Kyung Min Noh, Mikael Marttinen, Judith B. Zaugg

## Abstract

Enhancers and transcription factors (TFs) are crucial in regulating cellular processes, including disease-associated cell states. Current multiomic technologies to study these elements in gene regulatory mechanisms lack multiplexing capability and scalability. Here, we present SUM-seq, a cost-effective, scalable **S**ingle-cell **U**ltra-high-throughput **M**ultiomic sequencing method for co-assaying chromatin accessibility and gene expression in single nuclei. SUM-seq enables profiling hundreds of samples at the million cell scale and outperforms current high-throughput single-cell methods. We applied SUM-seq to dissect the gene regulatory mechanisms governing macrophage polarization and explored their link to traits from genome-wide association studies (GWAS). Our analyses confirmed known TFs orchestrating M1 and M2 macrophage programs, unveiled key regulators, and demonstrated extensive enhancer rewiring. Integration with GWAS data further pinpointed the impact of specific TFs on a set of immune traits. Notably, inferred enhancers regulated by the STAT1/STAT2/IRF9 (ISGF3) complex were enriched for genetic variants associated with Crohn’s disease, ulcerative colitis and multiple sclerosis, and their target genes included known drug targets. This highlights the potential of SUM-seq for dissecting molecular disease mechanisms. SUM-seq offers a cost-effective, scalable solution for ultra-high-throughput single-cell multiomic sequencing, excelling in unraveling complex gene regulatory networks in cell differentiation, responses to perturbations, and disease studies.

## Introduction

Gene regulatory elements in enhancer regions play a major role in disease development, as evidenced by the fact that most genome-wide association studies (GWAS) single nucleotide polymorphisms (SNPs) fall in non-coding gene regulatory regions (reviewed in^1^). Moreover, many diseases manifest through an imbalance of cellular differentiation, highlighting the importance of studying enhancer dynamics in the context of cellular differentiation. For example, many autoimmune diseases are characterized by an imbalance of pro- and anti-inflammatory immune cells^2–4^. Thus, studying such disease mechanisms requires a technology that allows the joint analysis of enhancer and transcription factor (TF) dynamics along a differentiation time course at single-cell resolution.

Recent developments in scalability of single-cell omics technologies, specifically single-cell RNA-seq (scRNA-seq) and single-nucleus (sn)ATAC-seq have revolutionized our understanding of the diversity of cell states and cellular responses to perturbations (reviewed in^5^). Particularly, multimodal profiling has enhanced our ability to unravel gene regulatory dynamics governing fundamental biological processes^6–8^. However, while scalability for individual modalities has become available (e.g., scifi-RNA-seq^9^, sci-RNA-seq^10^, dsci-ATAC-seq^11^, sci-ATAC-seq^12^), the majority of current multiomic methods are limited in scalability, multiplexing capability, or cost effectiveness (e.g., 10x multiome, ISSAAC-seq^13^), with a handful providing scalability in terms of number of cells assayed (e.g., SHARE-seq^14^, Paired-seq^15^), but at the expense of complexity (**Table 1**).

**Table 1:** Comparison between SUM-seq and other ATAC+RNA single cell methods.

| Method | Microfluidics/<br>plate-based | Modalities | Multiplexing | Costs | No° cells | Sensitivity |
| --- | --- | --- | --- | --- | --- | --- |
| **SUM-seq** | mf | ATAC/RNA | unlimited | \$ | ultra-high | high |
| **10x multiome** | mf | ATAC/RNA | no | \$\$\$\$ | high | high |
| **SNARE-seq2** | mf | ATAC/RNA | no | \$\$ | low | high |
| **Paired-seq** | mf | ATAC/RNA | no | \$\$ | ultra-high | low |
| **ASTAR-seq** | mf | ATAC/RNA | no | \$\$ | low | high |
| **ISSAAC-seq** | mf | ATAC/RNA | no | \$\$\$ | high | high |
| **Smart3-ATAC** | plate | RNA | yes-plate | \$\$ | low | high |
| **SHARE-seq** | plate | ATAC/RNA | yes, but<br>limited to<br>first index | \$\$\$ | ultra-high | low |
| **scCAT-seq** | plate | ATAC/RNA | no | \$\$\$ | low | high |

mf = microfluidics

Here, we present SUM-seq (**S**ingle-cell **U**ltra-high-throughput **M**ultiomic sequencing): a cost effective and scalable sequencing technique for multiplexed multiomics profiling. SUM-seq enables simultaneous profiling of chromatin accessibility and gene expression in single nuclei at ultra-high-throughput scale (up to millions of cells and hundreds of samples^9^). To this end, we refined the two-step combinatorial indexing approach, originally introduced by Datlinger et al.^9^ for snRNA-seq, tailoring it to the multiomic setup. In brief, accessible chromatin and nuclear mRNA first receive a sample-specific index by transposition and reverse transcription, respectively. Subsequently, a second index is introduced using a droplet-based microfluidic platform (e.g. 10x Chromium). The dual indexing allows overloading of microfluidic droplets while retaining the capacity to assign matched RNA- and ATAC-seq reads to the same individual cell.

We demonstrate the application of SUM-seq in two independent experimental setups, a species-mixing benchmarking experiment and a macrophage M1 and M2 polarization time course experiment, exemplifying its flexibility to accommodate complex experimental setups. We use both modalities to resolve temporal patterns of gene regulation, and integrate with genetic evidence to link regulatory networks to disease. Additionally, we validate the feasibility of decoupling sample collection from library generation, rendering the method well-suited for extensive atlas projects, time course experiments, and other experimental designs that involve prolonged sample collection periods.

## Main

### Overview of the SUM-seq method

SUM-seq is built upon the concept of a recently developed combinatorial fluidic indexing scRNA-seq approach, scifi-RNA-seq^9^, and a low-throughput multiomic assay, ISSAAC-seq^13^. SUM-seq has 5 key steps: In the first step, nuclei are isolated and fixed with glyoxal, then split into equal bulk aliquots in a multi-well format. In the second and third steps, unique sample indices (one per well) are introduced for the ATAC and RNA modalities. For ATAC, accessible genomic regions are indexed by Tn5 loaded with barcoded oligos. For RNA, the mRNA molecules are indexed with barcoded oligo-dT primers via reverse transcription (**Figure 1a, detailed library structures in Extended Data Figure 1a,b, Supplementary Table 1**). In step four, samples are pooled for Tn5-based tagmentation of the cDNA-mRNA hybrids to introduce a primer binding site necessary for binding of the barcoded oligonucleotides in the microfluidic system. In step five, nuclei are overloaded onto the microfluidic system (e.g., 10x Chromium), facilitating the encapsulation of multiple nuclei in a single droplet. Within these droplets, fragments are barcoded with the microfluidic barcode, resulting in dual barcoding of fragments with both a sample index and a droplet barcode enabling downstream demultiplexing and assignment of sequencing reads to individual nuclei. This is followed by an initial amplification of both modalities after which the library is split into two equal proportions and subjected to modality-specific amplification. At this stage, a library index can be introduced to enable multiplexing libraries from independent experiments on the sequencer. snATAC and snRNA libraries are sequenced using standard Illumina sequencing primers. Additionally, we have developed a scalable and reproducible Snakemake pipeline for processing SUM-seq data (https://git.embl.de/grp-zaugg/SUMseq; available upon publication), in which reads are assigned to sample indices and further demultiplexed to single-cell resolution by the droplet barcode (**Figure 1b**). From here, reads are mapped and a gene expression matrix and tile matrix are generated for RNA and ATAC, respectively. Modalities can be matched based on the assigned sample index-cell barcode combinations. The pipeline provides reproducibility, high scalability and flexibility for execution on any system (local, cluster, cloud). In brief, SUM-seq offers a user-friendly and rapid workflow for ultra-high-throughput single-cell multiomic sequencing that doesn’t require specialized instrumentation and is compatible with a widely available microfluidic platform.

**Figure 1.**
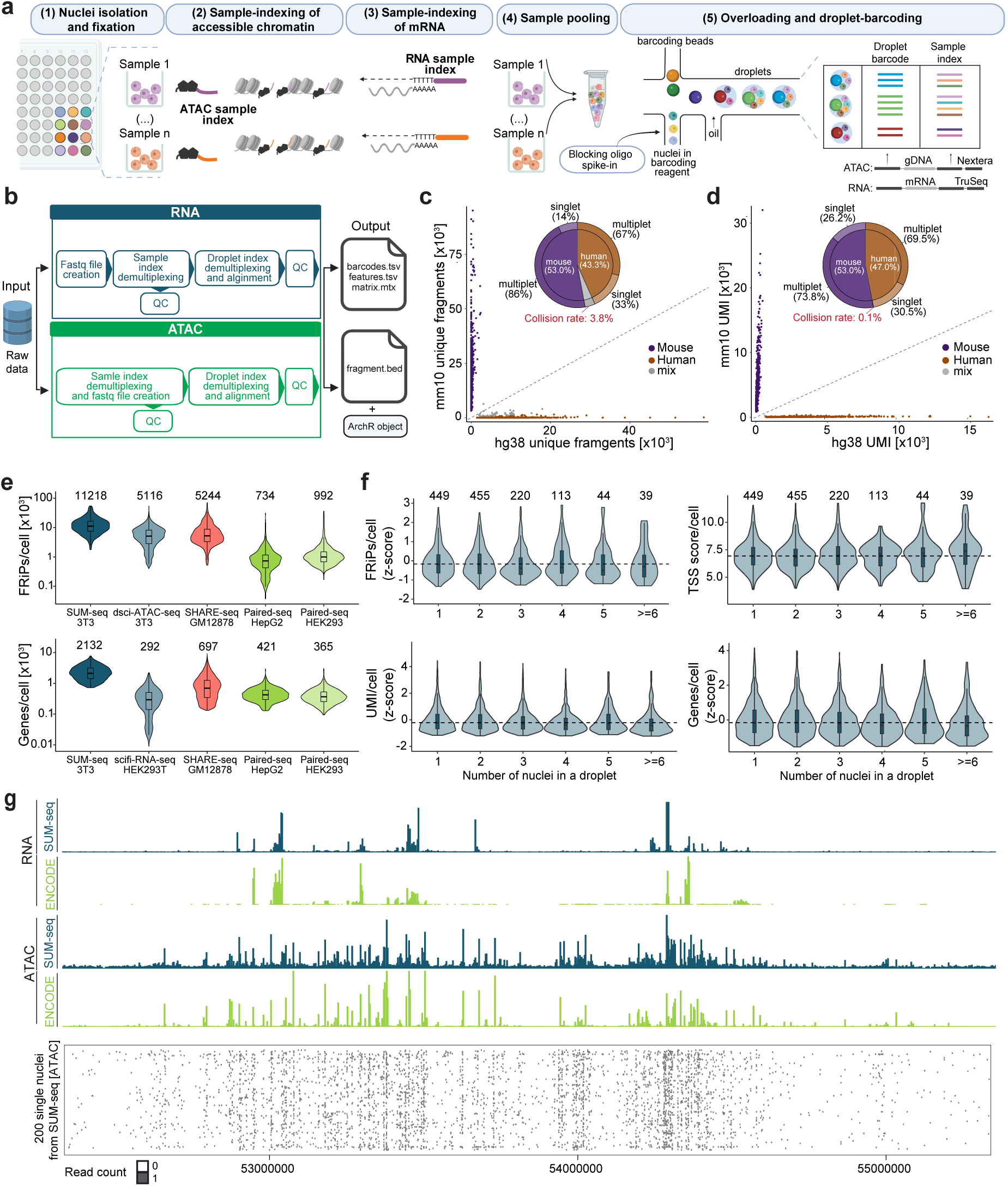
SUM-seq allows simultaneous profiling of chromatin accessibility and gene expression in single cells at ultra-high-throughput scale. **a,** Schematic depiction of the SUM-seq workflow. Key steps and detailed structures are described in the main text and **Extended Data Figure 1. b,** Schematic overview of the computational analysis pipeline. **c, d,** Species-mixing plots for the ATAC (**c**) and the RNA (**d**) modality, indicating the fraction of reads from singlets and multiplets assigned to the mouse genome (mm10, y-axes) and the human genome (hg38, x-axes). Genomic DNA fragments (**c**) and transcripts (**d**) are demultiplexed based on the combination of the sample index and the droplet barcode. The pie-chart shows the fraction of human and mouse cells as well as the frequency of multiplets and singlets. Collision rates are highlighted in red. **e,** Number of accessible DNA fragments in peaks (FRiP = fragments in peaks; upper panel) and genes detected (lower panel) per cell are shown as violin plots for SUM-seq and other methods (dsci-ATAC-seq^11^: GSM3507387; SHARE-seq^14^: GSM4156590 (ATAC), GSM4156602 and GSM4156603 (RNA); Paired-seq^15^: GSM3737488 (ATAC), GSM3737489 (RNA); scifi-RNA-seq^9^: GSM5151362). The median values are shown on top. **f,** Distribution of accessible DNA fragments in peaks per cell and the TSS score per cell (upper panels) as well as UMIs and genes detected per cell (lower panels) are shown as violin plots, split by the number of nuclei in a droplet. The number of droplets with N nuclei encapsulated is shown on the top. **g,** A representative genome browser view (*chr12*: *52,334,173 - 55,349,410*) of SUM-seq data is shown for K562 as aggregated transcript and ATAC fragment mappings. For comparison, K562 ATAC-seq and RNA-seq data tracks were retrieved from the ENCODE database (tracks labeled in green; GSE86660^16^ (RNA), GSE170214^16^ (ATAC)). The bottom panel shows binarised accessibility for 200 randomly selected K562 nuclei at single-cell resolution. Boxplots in **e,f**: center line represents median; lower and upper hinges represent the 25th and 75th quartiles respectively. The whiskers represent values that fall within 1.5 times the interquartile range between the 25th and 75th percentiles.

### SUM-seq enables efficient and scalable single-cell multiomics analysis

To evaluate the performance of SUM-seq and the data quality, we performed SUM-seq on an equal mixture of human leukemia (K562) and mouse fibroblast (NIH-3T3) cell lines. For this species-mixing benchmarking experiment, we loaded 100,000 nuclei into a single channel of the 10x Chromium system which equates to ∼7-fold overloading compared to the standard 10x workflow. To limit the sequencing requirements, we processed only 20 % (20,000 nuclei) of the generated microfluidic droplets for final library preparation. After combinatorial index demultiplexing, human and mouse reads were well-separated with a collision rate of 0.1 % (UMIs) and 3.8 % (ATAC fragments), resulting in 6,215 human cells and 7,607 mouse cells with data from both modalities (**Figure 1c**,**d**). This translates to a ∼70 % recovery rate of input loaded to the 10x Chromium system and represents a ∼7-fold increase in throughput at low collision rates compared to the standard 10x workflow (**Extended Data Figure 1d**). Key performance metrics of snRNA (UMIs and genes per cell) and snATAC (fragments in peaks per cell, TSS enrichment score, fragment size distribution) were consistently of very high quality for both modalities in SUM-seq and outperformed other high-throughput assays for scRNA, snATAC, and multiomic approaches (**Figure 1e, Extended Data Figure 2c-g**). Notably, there was no reduction in data quality for nuclei in overloaded droplets compared to droplets containing a single nucleus (**Figure 1f**). Lastly, the aggregate of snRNA and snATAC data resembled the published bulk RNA-seq and ATAC-seq in the K562 benchmarks from ENCODE data (**Figure 1g**; GSE86660^16^ (RNA), GSE170214^16^ (ATAC)).

During protocol optimization, we evaluated the effects of three factors on data quality: I) adding polyethylene glycol (PEG), II) methods to mitigate barcode hopping, and III) freezing of samples. (I) We found that, consistent with previous studies^17,18^, the addition of 12 % PEG to the reverse transcription reaction increased the number of UMIs and genes detected per cell (∼2.5- and ∼2-fold, respectively) with minor impact on the quality metrics of the ATAC modality (**Extended Data Figure 2h,i**). (II) We observed that barcode hopping can occur within multinucleated droplets^19,20^, primarily affecting the ATAC modality. However, we were able to mitigate it by using two complementary strategies: adding a blocking oligonucleotide^20^ in excess to the droplet barcoding step and reducing the number of linear amplification cycles during droplet barcoding from twelve to four (**Methods**) (**Figure 1b,c** and **Extended Data Figure 2a**). (III) To assess the suitability of SUM-seq for asynchronous sample collection (e.g. time course data or clinical samples), we tested glycerol-based cryopreservation following glyoxal fixation and found that it had minimal impact on the performance metrics of the assay (**Extended Data Figure 2j**). Thus, SUM-seq can be employed for gathering, fixing, and cryopreserving samples prior to library construction, which enables multiplexing samples in one experiment and thereby reducing costs and batch effects.

### SUM-seq recapitulates regulatory dynamics during M1/M2 macrophage polarization

SUM-seq is a suitable method for unraveling gene regulatory networks during cellular differentiation. As a model system for cellular polarization, we used macrophages, innate immune cells that can polarize towards a pro-inflammatory M1 or an anti-inflammatory M2 state depending on microenvironmental signaling. Despite the inherent complexity and heterogeneity observed *in vivo*, the M1/2 dichotomy serves as a useful framework for elucidating major transcriptional events and regulatory elements governing macrophage polarization. Previous research has determined TFs including STAT1, IRF5, and NF-κB as key factors involved in M1 polarization, and STAT6, PPARγ, and IRF4 in M2 polarization^21–23^. However, the complex regulatory networks orchestrating macrophage polarization over time remain elusive.

To unravel these intricate regulatory networks, we applied SUM-seq to profile hiPSC-derived macrophages across a polarization time course from the naive M0-state to the M1- and M2-states. We stimulated the M0 macrophages with LPS and IFN-γ to induce M1 polarization or IL-4 to induce M2 polarization. To discern early and sustained responses at chromatin accessibility and gene expression levels, we collected samples at five time points along the two polarization trajectories; prior to stimulation (M0) and at 1-hour, 6-hour, 10-hour, and 24-hour intervals, each sampled in duplicates totaling 18 samples **(Figure 2a)**. Leveraging the overloading capacity of SUM-seq, we loaded 150,000 nuclei into a single microfluidic channel of the 10x Chromium system, resulting in multiomic data from 51,750 high-quality nuclei evenly distributed across samples (**Figure 2b; Extended Data Figure 3**). For the snATAC, nuclei passing the QC filters (**Methods**) exhibited an average of 11,900 unique fragments, a TSS score of 8, and 40 % of reads in peaks. For the snRNA-seq readout, nuclei displayed on average 407 UMIs and 342 genes (**Extended Data Figure 3a-c**). The lower average UMI and gene counts per nucleus in the macrophage experiment, in comparison to the mixed-species experiment, are a consequence of the omission of PEG during reverse transcription due to a technical issue (**Methods**). However, the high number of nuclei that passed the quality filter enabled effective downstream analyses.

**Figure 2.**
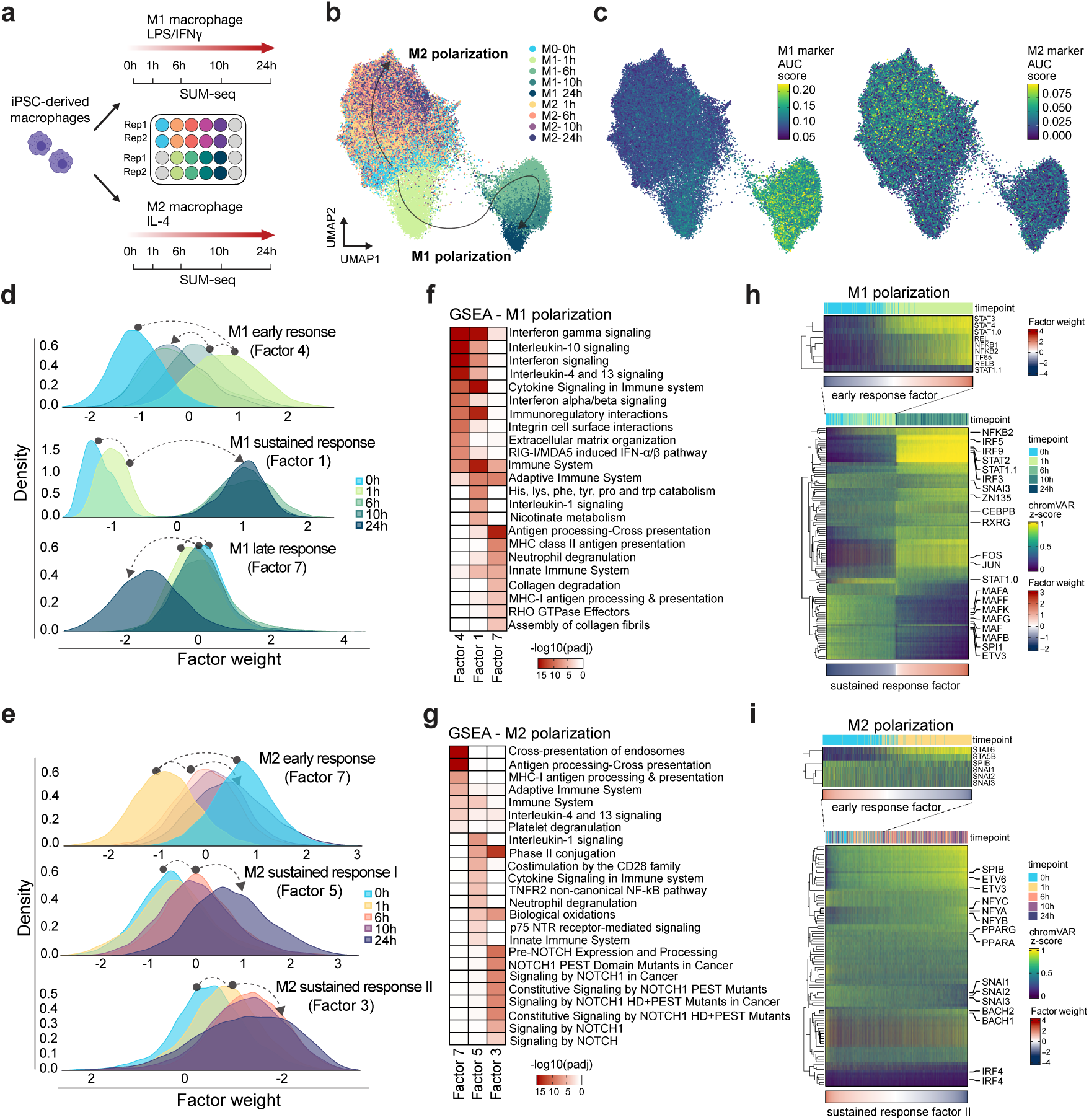
Integrated chromatin accessibility and gene expression data at single-nucleus resolution via SUM-seq characterizes differentiation trajectories along M1/M2 macrophage polarizations. **a,** Schematic overview of the macrophage polarization experiment. hiPSC-derived macrophages were stimulated with LPS and IFN-γ (M1) or IL-4 (M2). Nuclei were fixed with glyoxal and collected for SUM-seq at 0h, 1h, 6h, 10h, and 24h (*n*=2 per time point). **b,c,** Weighted nearest neighbor (WNN) UMAP projection of integrated SUM-seq data of macrophage polarization. Cells are annotated and labeled according to their sample index (**b**), AUC score of M1 signature genes (**c** left panel) and AUC score of M2 signature genes (**c** right panel; Supplementary Table 2). **d**,**e,** Distributions of cells from each time point across three MOFA factor weights associated with M1 polarization (**d;** early response, sustained response, late response) and M2 polarization (**e**; early response, sustained response I and II). Dotted arrows indicate the direction of the time-resolved response. **f**,**g,** Gene set enrichment analysis (GSEA) (Methods) for M1 (**f**) and M2 (**g**) polarization factors are shown as heatmaps. **h,** Motif activity for TFs associated with M1 early response (Methods) across M0 and M1-1h cells sorted by M1 early response factor weights (top). Motif activity for TFs associated with M1 sustained response across all M0 and M1 cells sorted by sustained factor weights (bottom). STAT1 and STAT1.1 represent the homodimer and heterodimer motifs of STAT1 respectively. **i,** Motif activity for TFs associated with M2 early response across M0 and M2-1h cells sorted by M2 early response factor weights (top). TF motif activity for TFs associated with M2 sustained response II across all M0 and M2 cells sorted by sustained factor II weights (bottom).

After preprocessing the data (**Methods**), we obtained a joint visualization of the two modalities by constructing a weighted nearest-neighbor (WNN) graph, which was further used to determine a uniform manifold approximation and projection (UMAP) (**Methods**). The resulting UMAP separated cells along the M1 and M2 polarization trajectories (**Figure 2b**). By calculating M1 and M2 scores from the expression of literature-based marker genes (**Supplementary Table 2; Methods**), and mapping them onto the UMAP, we observed increased M1 and M2 scores during the respective polarization trajectories (**Figure 2c**), confirming the validity of our experimental setup. M2 cells were not clearly distinguished from the M0 state and underwent only gradual changes throughout the time course. In contrast, during M1 polarization, we observed a clear separation of cells from the M0 state, with a substantial shift between the 1-hour and 6-hour time points. This separation of cells from different timepoints along the M1 and M2 trajectories underscores the capacity of our system to effectively capture distinct early and late polarization stages.

To delineate the key features that influence gene expression and chromatin accessibility during macrophage M1/M2 polarization, we performed Multi-Omics Factor Analysis (MOFA)^24^ to establish a shared low-dimensional representation for the snATAC- and snRNA-seq data (**Extended Data Figure 3d,e**). Investigating the association of the MOFA factors with metadata, we pinpointed three factors associated with M1 polarization, explaining the early response (Factor 4), sustained response (Factor 1), and late response (Factor 7) (**Figure 2d, Extended Data Figure 3d**). Notably, the early response was predominantly driven by chromatin accessibility, while the sustained and late responses were influenced by both gene expression and chromatin accessibility (**Extended Data Figure 3d**). For M2 polarization, we identified an early response (Factor 7) and two distinct sustained response factors (Factors 3 and 5) (**Figure 2e**). For functional interpretation of the MOFA factor associated genes, we performed gene set enrichment analysis (**Supplementary Table 3, Methods**). Genes associated with the early M1 response were enriched in IFN-γ and cytokine signaling (**Figure 2f**). IFN signaling was also enriched in genes associated with the sustained response (Factor 1), along with general immune system processes, IL-1 signaling, and metabolic terms, in line with metabolic reprogramming previously observed in M1 macrophages^25^. Genes associated with the M1 late response were predominantly enriched for terms related to antigen (cross-) presentation and RHO signaling (**Figure 2f**). The early M2 polarization factor was enriched for IL-4 signaling, while the two M2 sustained factors were enriched in biological oxidation - likely signifying the switch in metabolism of M2 macrophages^26^, and non-canonical NF-κB-signaling and Notch signaling (**Figure 2g**).

### SUM-seq uncovers key TF motif activity dynamics along the M1/M2 polarization

Next, we investigated which TFs underlie the chromatin remodeling during M1 polarization. Using chromVAR, we quantified TF motif accessibility variation^27^ and use this as a proxy for the activity of a TF binding a particular motif (here referred to as TF motif activity). We selected the top 10 % most variable TF motifs and filtered for those enriched in peaksets associated with either the early, the sustained, or the late response of M1 (**Extended Data Figure 4a**), resulting in 103 unique TFs (139 motifs; **Supplementary Table 4**).

When plotting the chromatin-based TF motif activity in cells from M0 and M1-1h along the early response (Factor 4) trajectory, we observed increasing motif activity for prototypical M1 TFs, such as STAT1 and IRF5^22,23^, followed by NF-κB (NFKB1, NFKB2, RELA, REL, TF65; **Figure 2h**). Furthermore, we observed a transient upregulation of the AP-1 complex motifs (JUN, FOS, etc.), indicative of a cellular activation state^28^. Meanwhile, the myeloid differentiation factors of the ETS family, including SPI1 (PU.1)^29^, were active in M0 macrophages and slightly decreased their activity upon polarization. To investigate the M1 response at later time points, we ordered all M1 cells along the sustained response factor (Factor 1; **Figure 2h**). This revealed a switch-like increase in motif activity for many prototypical M1 TFs, including STATs and IRFs, such as IRF5. STAT1 can function as a homodimer (also called IFN-γ activated factor (GAF)) or can form, together with STAT2 and IRF9, the interferon-stimulated gene factor-3 (ISGF3) complex^30^. STAT1 has two motifs in the HOCOMOCO database: one motif is bound by the STAT1 homodimer (STAT1.H12INVIVO.0.P.B), and the other motif (STAT1.H12INVIVO.1.P.B) is highly similar to the motif of STAT2. In the JASPAR database^31^, this second motif is classified as the interaction of STAT1 and STAT2. As such, herein, we refer to the second motif as the STAT1 heterodimer motif. Notably, we observed a steady drop in STAT1 homodimer motif activity over time, which coincided with a marked increase of its heterodimer motif activity as well as the motifs of STAT2 and IRF9 (**Figure 2h**). These observations highlight the role of STAT1 homodimer in initiating chromatin remodeling in response to IFN-γ. Meanwhile, the sustained response to IFN-γ is maintained by partnering with STAT2 and IRF9 in the ISGF3 complex, the driver of type-I IFN response^30^. Additional factors like AP-1 and ETS family members also exhibited switch-like activity patterns, increasing and decreasing with time, respectively. In contrast to the switch-like dynamic of the IRFs, STATs, AP-1 and ETS, the NFKB2 motif activity presented a more gradual increase along the M1 polarization. In the late response phase we observed only subtle changes in TF motif activity, including a slight increase in NFY motifs and a slight decrease in CEBP motifs (**Extended Data Figure 4c**). This pattern suggests that TF-driven gene regulation is stable between 6 and 24 hours of M1 polarization.

A similar analysis of TFs along the M2 polarization focused on 93 TFs (121 motifs) (**Extended Data Figure 4b, Supplementary Table 4**). Overall, we observed less striking dynamics along the M2 polarization trajectories compared to the M1 response. The most notable pattern was observed for the IL-4-responsive STAT6^32^, which showed increasing motif activity along the early M2 response trajectory (**Figure 2i**). The sustained response patterns were mostly characterized by a steady increase in motif activity for ETS family members, most notably SPIB^33^, ETV transcription factors, such as ETV3 and ETV6^34^, and increased activity of NFY factors (**Figure 2i, Extended Data Figure 4d**). Conversely, SNAIL family member motif activity decreased along the sustained response. Likewise, accessibility of CTCF, a protein shaping chromatin structure^35^, decreased upon M2 polarization, suggesting a global rewiring of chromatin architecture.

Overall, we observed strong remodeling of the chromatin landscape along the macrophage polarization trajectories, much of it driven by specific sets of TFs.

### Gene regulatory network analysis reveals sustained response by STAT1/STAT2/IRF9 in M1 macrophages

We next aimed to understand how the TF-driven chromatin remodeling leads to activation of specific gene expression programs. To that end, we constructed an enhancer-mediated gene regulatory network (eGRN) using GRaNIE^36^ (**Supplementary Table 5; Methods**). Out of the 103 TFs identified in association with M1 polarization, we retrieved 44 within the eGRN, either as a TF itself or as a target gene of the M1-TFs (**Figure 3a; Extended Data Figure 5b**). The eGRN shows that the most connected TFs share many of their target genes. For example, IRF1 and IRF8, connected to 1002 and 804 genes respectively, share 728 of their regulon genes.

**Figure 3.**
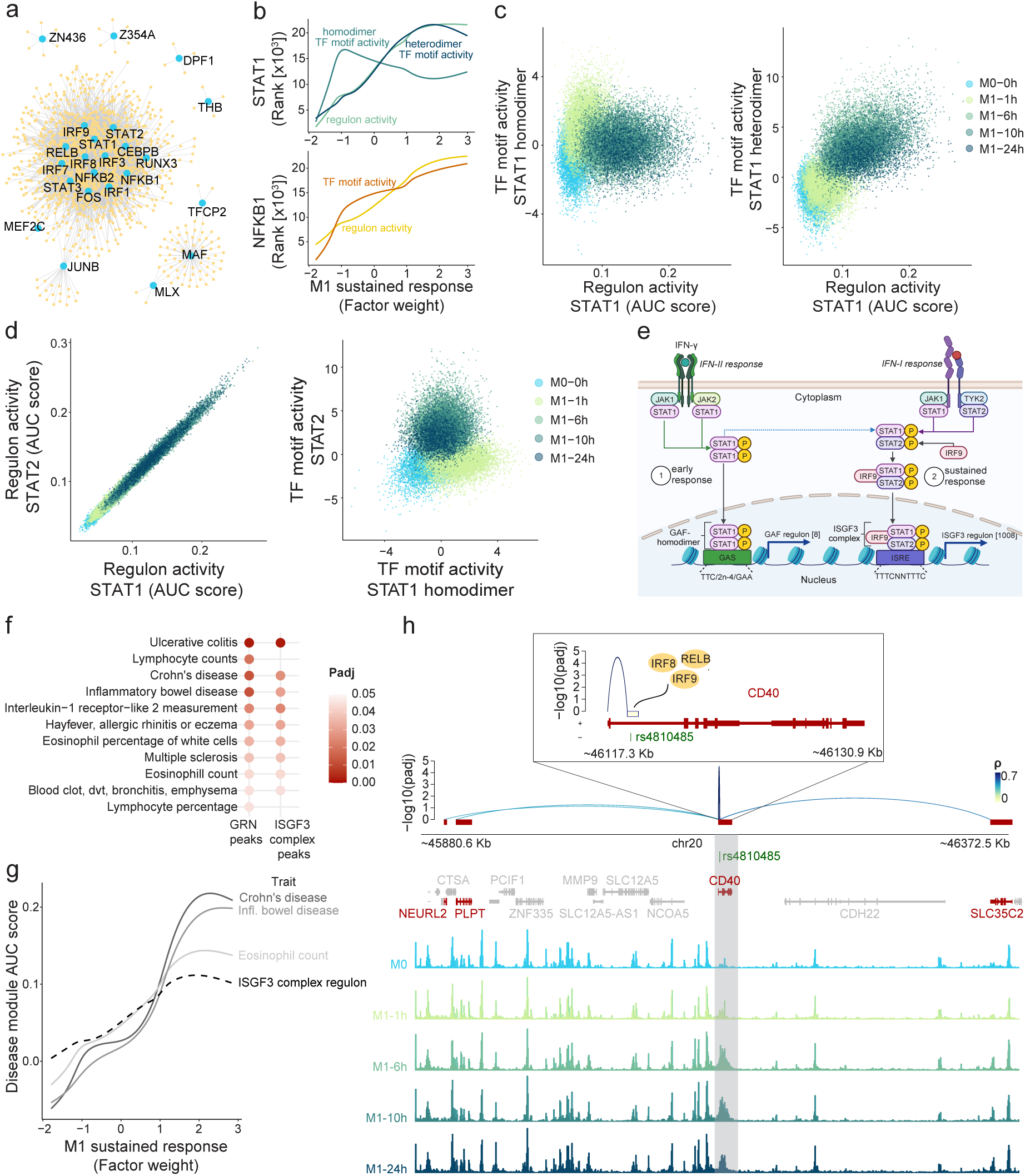
Gene regulatory networks inferred from SUM-seq data in macrophages coupled with genetic evidence reveals TF and regulon hierarchy, linking TFs to immune traits. **a,** Gene regulatory network visualization of TFs and their assigned target genes, including only TFs with at least 4 target genes. **b,** TF motif activity (chromVAR Z-score) and regulon activity (AUC cell score) for STAT1 (top) and NFKB1 (bottom) motifs along the M1 sustained response factor. Cells are ranked by their respective value for activity measures, lines show best fit (generalized additive model). **c,** Scatter plots of the correlation between the STAT1 regulon activity and STAT1 homodimer (left) and STAT1 heterodimer (right) TF motif activity (chromVAR Z-score). Points represent cells and are coloured by their experimental time point. **d,** Scatter plots of each cell across the M1 macrophage polarization showing the correlation between STAT1 and STAT2 regulons (left) and STAT1 homodimer and STAT2 TF motif activity (right). Points represent cells and are coloured by their experimental time point. **e,** Schematic of the type II IFN-associated STAT1 homodimer (known as GAF) response, representing the early response, and the type I IFN-associated STAT1/STAT2/IRF9 (known as ISGF3 complex), representing the sustained response. IFN-γ binding to its receptor initiates an early response characterized by the phosphorylation and subsequent dimerization of STAT1, forming GAF. The GAF-homodimer then translocates into the nucleus and binds the GAS motif, activating the expression of 8 downstream targets identified in our eGRN. As a secondary response, STAT1, STAT2 and IRF9 undergo phosphorylation and activation, forming the ISGF3 complex. The ISGF3 complex binds the ISRE motif, potentially activating the expression of 1008 downstream genes identified in our eGRN. **f,** Identification of diseases and traits associated with genetic variants enriched in open regions of all GRN peaks and a highly connected regulon in the eGRN (the STAT1/STAT2/IRF9 - putative ISGF3 complex) using linkage disequilibrium score regression (LDSC). All open regions in macrophages are used as background. **g,** AUC cell scores for disease modules derived by intersection of putative genome-wide SNPs for the top enriched diseases with the putative ISGF3 complex regulon (union of STAT1, STAT2, IRF9 gene targets) (**Methods**), compared to AUC cell scores for all ISGF3 regulon genes along the M1 sustained response factor. **h,** eGRN peak-gene interactions for the CD40 intronic peak intersecting RA-associated SNP rs4810485 (top), zooming in to the CD40 region with the TFs binding to this peak (insert). M0 and M1 cell aggregate ATAC-seq tracks split by timepoint highlighting the CD40 intronic peak (bottom).

We further resolved the hierarchy of a TF’s activity on chromatin accessibility and their effect on inducing transcription by comparing motif and regulon activity similarity along the M1 immediate response factor (**Methods**). For the majority of TFs, we found a positive correlation between their motif activity and the expression of their target genes derived from the eGRN (**Extended Data Figure 6a**). Some evident exceptions to this were STAT1, STAT3, NFKB1, NFKB2 and RELB. For NFKB1, we observed a slightly delayed expression of its regulon despite the gradual increase in NFKB1 motif activity from an early time point (**Figure 3b**). Additionally, we found that STAT1 regulon activity was discordant with the chromatin activity for the STAT1 homodimer motif, but it closely tracked the STAT1 heterodimer motif activity (**Figure 3b,c**).

While the STAT1 homodimer motif activity was strongest at 1h post-stimulation and decreased from 6h onwards, the STAT1 heterodimer motif, along with STAT2 and IRF9 motif activities, exhibited a gradual increase over time (**Figure 3d, Extended Data Figure 6b,c**). This pattern was also reflected by the STAT1, STAT2 and IRF9 regulons. These three regulons shared 601 genes and their activities were highly correlated, providing evidence that the ISGF3 complex was captured by the eGRN (**Figure 3d**; **Extended Data Figure 6d-f**). In contrast, we identified only 8 target genes connected to the homodimer motif, all of which were also connected with the heterodimer motif (**Supplementary Table 6**). After IFN-γ/LPS stimulation, we first observe increased activity for the STAT1 homodimer and other early response TFs (**Figure 2h**). This early response potentially leads to endogenous type I IFN production, which in turn activates the ISGF3 complex, explaining the delayed response of ISGF3 as compared to the STAT1 homodimer activity (**Figure 3e**).

Given that the activity of many TFs, including STAT1, is regulated by phosphorylation, we made use of published phosphoproteomic data on IFN-γ/LPS and IL-4 stimulated THP1-derived macrophages to validate our findings^37^. We observed that upon IFN-γ/LPS stimulation, STAT1 becomes phosphorylated at sites T699 and Y701 almost immediately, followed by a gradual decrease over time (**Extended Data Figure 6g**). Phosphorylation of these sites is known to be induced by IFN signaling and triggers STAT1 accumulation in the nucleus and activation of DNA binding activity^38,39^. This pattern thus recapitulates the early STAT1 homodimer response we observed on chromatin. Moreover, the observed gradual increase in STAT2 and IRF9 activity based on ATAC-seq and RNA-seq data is corroborated by a delayed but sustained increase in the phosphorylation of IRF9 at sites S131 and S253 (**Extended Data Figure 6h**). S253 is known to be induced by IFN-β (a type I IFN) and may play a role in regulating the expression of interferon-stimulated genes and interactions with STAT2^40^.

In summary, integrating chromatin accessibility and gene expression data at single-cell resolution over a time course, we characterized the M1 and M2 polarization trajectories. This highlights SUM-seq’s capability to provide detailed insights into cellular states and TF activities. Using both ATAC and RNA modalities along a time course enabled us to dissect the regulatory layers of M1 polarization, leading to the identification of a putative hierarchy of IFN-driven responses. Specifically, we observed a marked shift in STAT1-mediated regulation, from its homodimer-driven chromatin remodeling during the early M1 polarization, to an ISGF3-driven response reflected in both accessibility and transcription at later time points. This switch is in line with previous reports on the timing of IFN stimulations^41,42^ (**Figure 3e**).

### GWAS enrichment analyses of the SUM-seq identified eGRN

To further investigate the role of the macrophage regulatory network in disease, we tested whether the chromatin accessible regions linked to regulon genes in the network were enriched for heritability of specific diseases using linkage disequilibrium score regression (LDSC)^43^.

Overall, we found all open regions in our macrophage dataset strongly enriched for white blood cell count traits and autoimmune diseases (**Extended Data Figure 7a**). Therefore, we used all open regions as background, to ensure the disease enrichments are not driven by the cell type only. The eGRN peaks were enriched for specific immune-related traits including inflammatory bowel disease (IBD), two subtypes of IBD (ulcerative colitis (UC) and Crohn’s disease (CD)), hayfever and multiple sclerosis (**Figure 3f**). Mapping the putative genome-wide SNPs (p < 10^-6^) for the top heritability-enriched diseases and traits (p < 0.1), we found 112 unique SNPs that directly overlap with the peaks in the eGRN (**Supplementary Table 7**). For UC, the most significant overlapping SNP was rs153109, located in an intron of the Interleukin-27 (IL-27) gene (**Extended Data Figure 7b**). This SNP has also been associated with CD and with IBD. These diseases are characterized by chronic inflammation of the digestive tract. IL-27 is mainly produced by antigen presenting cells including macrophages, and has been implicated in the regulation of the immunological response in IBD^44^. As such, it has been proposed as a potential new drug target for IBD^45^. In our eGRN, this IL-27 peak is connected to and putatively regulated by NFKB1. Interestingly, a NFKB1 knockout in a THP1 monocyte cell line showed downregulation of IL-27, providing further evidence for its regulatory function^46^.

To zoom in further, we tested heritability enrichment for the peaks in the putative ISGF3 regulon (union of peaks linked to STAT1, STAT2 and IRF9). Although this subset consisted of only half as many peaks as the full eGRN (1673 vs. 3217), the enrichments for the autoimmune diseases remained significant (**Figure 3f**). This suggests that STAT1, STAT2 and IRF9 play an important role in autoimmune diseases involving macrophages. Indeed, disease SNPs mapping specifically to the ISGF3 regulon resulted in prioritizing 19 genes for IBD, 16 for CD, and 37 for eosinophil counts (**Supplementary Table 8; Methods**). We then calculated the expression activity for these disease genes (**Methods**) and projected them along the M1 sustained response factor. The disease gene sets (IBD, CD) showed a stronger increase in activity over time than the full ISGF3 regulon and “eosinophil count” gene set, suggesting the polarized M1 state is particularly important for these diseases and traits (**Figure 3g**). Another example bridging the M1 polarization and genetics is rs4810485, which overlaps with the ISGF3 regulon. This SNP, located in the intron of CD40, is associated with IBD, CD and UC and the CD40 intronic peak is linked to CD40, PLTP, NEURL2 and SLC35C2 in our eGRN (**Figure 3h**). Previous work has linked this SNP to CD40 as an expression, protein and splice quantitative trait locus (e-, p-, sQTL) in blood and monocyte-specific datasets, as well as to PLTP as a blood eQTL^47^ (**Supplementary Table 9)**. CD40 is a cell surface receptor expressed by antigen presenting cells including macrophages, that contributes to the activation of T-cells through its interaction with CD40 ligand (CD40-L). Given this role, CD40 is likely involved in the pathogenesis of IBD and other autoimmune diseases^48^. CD40 expression can be induced by IFN-γ stimulation in macrophages, a process thought to be involved in the initiation of chronic inflammation in IBD^49,50^.

In summary, the integration of our high-resolution single-cell multimodal time course data on M1 polarization with genetic disease evidence revealed molecular disease mechanisms and suggests novel candidate pathways that will require validation in future studies.

## Discussion

SUM-seq is a single-cell multiomic method that allows highly multiplexed and scalable experimental setups to investigate the molecular underpinnings of gene regulation via profiling of chromatin accessibility and gene expression from the same cell. In this study, we applied SUM-seq to examine macrophage polarization, demonstrating how the multimodal data from a single multiplexed experiment can unravel the molecular underpinnings of TF-driven disease mechanisms, integrating TF response dynamics with genetic evidence from GWAS.

We benchmarked SUM-seq, showcasing its ability to effectively capture both chromatin accessibility and gene expression at the singel-cell level, outperforming previously published scalable high-throughput methods. Notably, we demonstrated its capability to generate high-quality data from both fresh and cryopreserved samples, making it adaptable for extensive atlas studies, where sample collection can span a long time period and requires multi-center efforts. While SUM-seq does not reach the gene expression complexity of lower throughput droplet-based single-cell multiomic methods, it provides matching chromatin accessibility complexity paired with significant improvement in cost-efficient scalability. We anticipate that through minor optimisations 1) gene expression complexity will further improve and 2) SUM-seq can expand to incorporate surface proteomic readouts by use of single cells instead of single nuclei^51^. Furthermore, it should be noted that although the current format measures chromatin accessibility and gene expression, the strategy can be adopted for the measurement of other omic layers such as TF binding and histone modifications (Cut&Tag^52^), DNA methylation (sci-MET^53^), and whole genome sequencing (s3-WGS^54^).

We applied SUM-seq to hiPSC-derived macrophages in a time-course of M1 and M2 polarization, as a prototypical model of cellular polarization. By analyzing the activity of TF motifs on the chromatin versus the expression of associated TF target genes, we observed a hierarchical activation of a TF cascade upon IFN-γ/LPS stimulation of M0 macrophages. Specifically, we identified TFs that clearly act in a stepwise manner (i.e. STAT1), whereas others exhibited a continuous activation profile (i.e. NFKB1). We could recapitulate the hierarchy of STAT1 homodimer activity preceding STAT1/STAT2/IRF9 activity. A potential explanation for the delay in the activity of the NFKB regulon with respect to its motif activity is the previously reported observation that enhancers of NF-κB response genes are primed by NF-κB but their expression additionally requires activity of the IFN-I-induced ISGF3 complex^55^.

Genes with genetic links to disease have been shown to be better drug targets^56^. Yet for the majority of disease-associated genetic variants, we lack molecular mechanisms and it is often unclear in which cell type or disease state a variant exerts its effect. Most disease-associated common genetic variants lie in non-coding regulatory elements that are presumably regulated by TFs. Indeed, TFs are often identified as main drivers of disease-relevant phenotypes in genetic screens^57^. Our study emphasizes that understanding disease mechanisms requires integrating genetic evidence with disease-specific gene-regulatory dynamics. SUM-seq was instrumental in pinpointing gene-regulatory dynamics and linking them to genetic disease evidence in a single experiment. Specifically, profiling the macrophage polarization response in a single SUM-seq experiment, followed by integrative analysis of the eGRN and publicly available summary statistics from GWAS, identified several known disease mechanisms, as well as novel associations. For example, our findings reinforce the link between the ISGF3 complex, CD40 expression, and IBD. Indeed, the role of macrophages in maintaining intestinal immune homeostasis makes them an appealing target to develop new therapeutic strategies for IBD^58^.

In conclusion, our study showcases the effective implementation of SUM-seq in the context of a time course experiment. Importantly, our findings suggest broad applicability of this method to diverse experimental settings that demand gene regulatory analyses at single-cell resolution. Specifically, SUM-seq is a promising tool for arrayed CRISPR, drug or perturbation screens, and large-scale atlas projects, underscoring its potential as a versatile and powerful technique across a spectrum of biological studies. Overall, we envision SUM-seq as an easily adoptable, scalable method for projects requiring multiomic single-cell profiling up to millions of cells from hundreds of samples.

## Methods

### Cell culture

Human myelogenous leukemia cells (K562, DSMZ no. ACC 10) were cultured in RPMI 1640 medium (Gibco, catalog no. 11875093) supplemented with 10 % FBS (Gibco, catalog no. 10270-106), 100 U/ml penicillin/streptomycin (PenStrep, Gibco, catalog no. 15140122) and 1x non-essential amino acids (Gibco, catalog no. 11140050) at 37°C with 5 % CO2. Mouse fibroblast cells (NIH-3T3, DSMZ no. ACC 59) were cultured in DMEM medium (high glucose, Gibco, catalog no. 11965092) supplemented with 10 % FBS, 100 U/ml penicillin/streptomycin and 1x non-essential amino acids at 37 °C with 5 % CO2.

### Macrophage differentiation

hiPSCs (CESCG-295; male, peripheral blood mononuclear cell-derived; provider: Dr. Michael Snyder, Stanford University; approved Material Transfer agreement) were cultured in Essential 8 medium (Gibco, catalog no. A1517001) in dishes coated with vitronectin (ThermoFisher, catalog no. A14700). hiPSCs were differentiated to macrophages as previously described by van Wilgenburg et al^59^. In brief, 4 million hiPSCs were resuspended in EB medium (Essential 8 medium (Gibco, catalog no. A1517001), 50 ng/ml Recombinant Human BMP4 (Peprotech, catalog no. 120-05ET), 50 ng/ml Recombinant Human VEGF (Peprotech, catalog no. 100-20), 50 ng/ml Recombinant Human SCF (Peprotech, catalog no. 300-07)) with 10 μM Y-27632 (AbCam Biochemicals, catalog no. ab120129), seeded in 400-microwell Aggrewells (Stemcell Technologies, catalog no. 34450) and centrifuged at 100 x g for 3 min to evenly distribute the cells in microwells. 75 % of the medium was replaced by fresh EB medium on the next 2 consecutive days. On the third day, EBs were transferred to a low-attachment plate (Sigma-Aldrich, catalog no. CLS3471-24EA) and cultured for two additional days with minimal disruption. On the fifth day, EBs were transferred to the final format of choice and left undisturbed for 1 week in factory media (*X-VIVO*^TM^-15 (Lonza, catalog no. BE02-060F) 1 % GlutaMax (Thermo Scientific, catalog no. 35050061), 1 % PenStrep (Gibco, catalog no. 15140122), 50 µM ß-mercaptoethanol, 50 µg/ml Normocin (Invivogen, catalog no. ant-nr-05) and 100 ng/ml Recombinant Human M-CSF (Peprotech, catalog no. 300-25), 25 ng/ml Recombinant Human IL-3 (Peprotech, catalog no. 200-03)) to allow them to attach. Fresh media was added weekly until macrophage precursor production started around week 4, as identified by the presence of large suspension cells with spherical morphology. From that point on, factories were maintained for a total of 8-10 weeks. Precursors were harvested weekly, taking approximately 1/4^th^ of the total volume containing precursors and replacing it with fresh media. For the terminal differentiation of precursors to macrophages, harvested precursors were resuspended in macrophage media (*X-VIVO*^TM^-15, 1 % GlutaMax, 1 % PenStrep, Recombinant Human M-CSF) and plated in the desired format at a density of 160.000 cells/cm^2^ for 7 days.

### Polarization to M1 or M2 macrophages

At day 7, mature macrophages were polarized to M1 or M2. For M1 polarization, macrophages were cultured in macrophage media supplemented with 20 ng/ml human recombinant IFN-γ (PeproTech, catalog no. 300-02) and 25 ng/ml LPS (Invivogen, catalog no. tlrl-3pelps). For M2 polarization macrophages were cultured in macrophage media supplemented with 20 ng/ml recombinant human IL-4 (PeproTech, catalog no. 200-04). Cells were harvested after 24, 10, 6 and 1h, including an undifferentiated control (M0) in which medium was replaced by fresh macrophage medium.

### Nuclei preparation

For the species mixing experiments, mouse NIH-3T3 cells were washed once with PBS and dissociated with Accutase (Stem Cell Technologies, catalog no. 07920). Two million NIH-3T3 and K562 cells were collected and fixed in a 3 % glyoxal solution (40 % glyoxal (Merck, catalog no. 128465), 0.75 % acetic acid; adjusted to pH 5 by addition of 1 M NaOH) for 7 min at room temperature. For cryopreservation, cells were washed after fixation with RSB-1%BSA-RI (10 mM Tris-HCl pH 7.5, 10 mM NaCl, 3 mM MgCl_2_, 1 mM DTT, 1 % BSA, 20 µg/ml in-house produced RNasin (referred to RNasin hereafter)) and slowly frozen (Freezing buffer: 50 mM Tris-HCl pH 7.5, 5 mM MgAc, 0.1 mM EDTA and 25 % glycerol). On the experiment day, fresh cells were processed as described above, while cryopreserved cells were thawed slowly on ice and washed with RSB-1%BSA-RI. Nuclei were extracted with 1x lysis buffer (10 mM Tris-HCl pH 7.5, 10 mM NaCl, 3 mM MgCl2, 0.1 % Tween20, 20 µg/ml RNasin, 1 mM DTT, 1 % BSA, 0.025 % IGEPAL CA-630 and 0.01 % Digitonin) incubating samples for 5 min on ice. Next, nuclei were washed with wash buffer (10 mM Tris-HCl pH 7.5, 10 mM NaCl, 3 mM MgCl_2_, 0.1 % Tween20, 20 µg/ml RNasin, 1 mM DTT, 1 % BSA) and filtered through a 40 µm cell strainer (Falcon, catalog no. 352340).

After the desired polarization time, macrophages were harvested by replacing the medium with ice-cold PBS-1%BSA and 2 mM EDTA and pipetting. Nuclei were extracted by resuspending the cell pellet in 1x lysis buffer for 4 min on ice and washed with wash buffer. To maintain nuclei integrity, nuclei were fixed with a 3 % glyoxal solution for 7 min at room temperature and washed with RSB-1%BSA-RI.

### Transposome generation

Tn5 was produced in-house according to a previously described protocol with minor modifications. In short, the pETM11-Sumo3-Tn5(E54K,L372P) plasmid was transformed into *Escherichia coli* BL21(DE3) codon + RIL cells (Stratagene). Cells were grown in TB-FB media (1.2 % Bacto-Tryptone, 2.4 % yeast-extract, 0.4 % glycerol, 17 mM KH_2_PO_4_, 72 mM K_2_HPO_4_, 1.5 % Lactose, 0.05 % Glucose, 2 mM MgSO_4_) supplemented with kanamycin and chloramphenicol at 37 °C until OD600 ∼0.5. The temperature was then lowered to 18 °C and cells were grown overnight, wherafter cells were harvested by centrifugation. The cell pellet was resuspended in running buffer (20 mM HEPES-NaOH pH 7.2, 800 mM NaCl, 5 mM imidazole, 1 mM EDTA, 2 mM DTT, and 10 % glycerol) supplemented with cOmplete protease inhibitors (Roche) and lysed using a Microfluidizer. Polyethyleneimine (PEI) pH 7.2 was added dropwise to a final concentration of 0.5 % to remove nucleic acids. The cleared lysate was loaded onto a cOmplete His-tag purification column (Roche) and the His6-Sumo3-Tn5 was eluted with a running buffer containing 300 mM imidazole. To remove the fusion tag, His6-tagged SenP2 protease was added to the elution fractions. The sample was digested overnight at 4 °C while being dialyzed back to the running buffer. The next morning, the dialyzed sample was loaded again onto a cOmplete His-tag purification column (Roche), and the untagged Tn5 was collected in the flow-through. The concentrated flow through was then loaded onto a HiLoad Superdex200 16/600 pg column (GE Healthcare) equilibrated with 50 mM Tris pH 7.5, 800 mM NaCl, 0.2 mM EDTA, 2 mM DTT, and 10 % glycerol. Elution fractions corresponding to the Tn5 dimer peak were pooled, aliquoted, snap frozen in liquid nitrogen and stored at -80 °C. For transposome assembly, 100 µM of oligonucleotides with the Nextera Read1 sequence and 100 µM of oligonucleotides with a Read2-sample_index-spacer structure were annealed with 100 µM mosaic end-complement oligonucleotides with a 3’ dideoxynucleotide end (ddc) at a 1:1:2 ratio by heating for 95 °C for 3 minutes followed by cooling down to 25 °C at a ramp rate of -1 °C/min. Annealed oligos were mixed with an equal volume of 100 % glycerol and stored at -20 °C until use. Finally, the annealed oligos were mixed with the in-house produced Tn5 (at 1 mg/ml in 50 mM Tris, 100 mM NaCl, 0.1 mM EDTA, 1 mM DTT, 0.1 % NP-40, and 50 % glycerol), at a 1:1 ratio and incubated for 30-60 min at room temperature. For the species-mixing experiment, annealed oligos were first diluted with H_2_O at a 1:1 ratio before generating the transposomes. Assembled Tn5 was stored at -20 °C until use.

### SUM-seq procedure

#### DNA transposition

For accessible DNA transposition, isolated and fixed nuclei were resuspended in a transposition mix containing 38.8 mM Tris-acetate, 77.6 mM potassium acetate, 11.8 mM magnesium acetate, and 18.8 % dimethylformamide supplemented with 0.005x protease inhibitor cocktail (Roche, catalog no. 11697498001), 0.4 U/µl SUPERaseIn (Thermo Fisher, catalog no. AM2694), 1.2 U/µl RiboLock (Thermo Fisher, catalog no. EO0382) and the barcoded Tn5 transposomes (1:10 (v/v); 938 µM) and incubated at 30 °C for 30 min with shaking at 400 rpm (n=2 per condition). The reaction was terminated by adding an equal volume of 2x stop buffer (10 mM Tris-HCl pH 7.5, 20 mM EDTA pH 8.0, 2 % BSA).

#### *In situ* reverse transcription

Transposed nuclei were washed 3 times with RSB-1%BSA-RI and resuspended in reverse transcription mix containing 1x RT buffer (50 mM Tris-HCl pH 8.0, 75 mM NaCl, 3 mM MgCl_2_, 10 mM DTT), 0.5 mM dNTPs, 1 U/µl Protector RNase inhibitor (Roche, catalog no. 3335402001), 0.2 U/µl SUPERaseIn (Thermo Fisher, catalog no. AM2694), 10 U/µl Maxima H Minus RT (Thermo Fisher catalog no. EP0752) and 5 µM of the respective barcoded oligo-dT primer (n=2 per condition). In the NIH-3T3/K562 species mix experiment, 12 % PEG 8000 (w/v; Jena Bioscience, CSS-256) was included in the RT mix as a crowding agent. For the macrophage experiment, crowding agent PEG was omitted from the reverse transcription reaction as we observed formation of white precipitates, which hampered downstream steps of library preparation. This issue was however later resolved by omitting BSA from pre- and post reverse transcription washes. The samples were incubated at 50 °C for 10 min, followed by 3 thermal cycles (8 °C for 12 sec, 15 °C for 45 sec, 20 °C for 45 sec, 30 °C for 30 sec, 42 °C for 2 min and 50 °C for 3 min), and finally incubated at 50 °C for 5 min. Following the RT reaction, all samples were pooled in RSB-RI and washed twice with RSB-1%BSA-RI.

#### cDNA/mRNA hybrid tagmentation

For transposome preparation, 100 µM of oligonucleotides with a Nextera Read1 sequence overhang were annealed with 100 µM of the Tn5-mosaic end oligonucleotides at a 1:1 (v/v) ratio by heating for 95 °C for 3 min followed by cooling down to 25 °C at a ramp rate of -1°C/min. Annealed oligos were mixed with an equal volume of 100 % glycerol and stored at - 20 °C until use. For transposome assembly, annealed oligos were mixed with the in-house produced Tn5 at a 1:1 ratio for 30 min at RT and stored at -20 °C until use.

The collected nuclei pool was counted, and the reaction volume was adjusted according to the total number of nuclei (100 µl reaction for each 100,000 nuclei). Nuclei were resuspended in the transposition mix containing 1x transposition buffer (38.8 mM Tris-acetate, 77.6 mM potassium acetate, 11.8 mM magnesium acetate, and 18.8 % dimethylformamide and 0.005x protease inhibitor cocktail and a 1:100 ratio (v/v) of assembled Tn5 (93.8 µM). The tagmentation was performed on a shaking thermoblock at 37 °C for 30 min at 400 rpm. The reaction was terminated by adding an equal volume of 2x stop buffer. Next, the sample pool was washed twice with RSB-1%BSA.

#### Gap fill and ExoI

Sample pool was resuspended a mix containing 1x RT buffer (50 mM Tris-HCl pH 8.0, 75 mM NaCl, 3 mM MgCl_2_, 10 mM DTT), 0.5 mM dNTPs, 8 U/µl Maxima H Minus RT (Thermo Fisher catalog no. EP0752) and 2 U/µl Thermolabile Exonuclease I (New England Biolabs, catalog no. M0568S) and incubated for 15 min at 37 °C on a shaking thermoblock at 400 rpm. Afterwards the sample was washed twice with RSB-1%BSA filtered through a 40 µm cell strainer and count.

#### GEM generation

Complex samples with a high proportion of debris were cleaned prior loading of the 10x Genomics chip using a glycerol-based buffer (50 mM Tris-HCl pH 7.5, 5 mM Mg-acetate, 0.1 mM EDTA, 25 % glycerol). For this, nuclei were resuspended in a glycerol-based buffer, layered on top of an additional 500 µl glycerol-based buffer and centrifuged at 800 x g for 15 minutes. The supernatant was discarded, and the pellet was washed twice with RSB-1%BSA and filtered through a 40 µM Flowmi cell strainer (Sigma-Aldrich, catalog no. BAH136800040). The nuclei were counted and loaded into the Chromium Controller (10x Genomics) according to 10x Genomics single-cell-ATAC standard protocol. 100,000 nuclei were loaded for the NIH-3T3/K562 species-mixing experiment and 150,000 for the macrophages experiment. In brief, nuclei in 1x Nuclei buffer (10x Genomics) were mixed with barcoding reagents (Barcoding Reagent B, Reducing Reagent B and Barcoding Enzyme). For the NIH-3T3/K562 species mix experiment, this reaction mix was supplemented with ∼500 nM of a blocking oligonucleotide to prevent Tn5-barcode hopping in multinucleated droplets. In brief, barcode hopping may occur when residual barcoded Tn5 molecules, which contain the ATAC sample index, are carried over to the droplets. During droplet barcoding, these residual transposomes are disassembled, and barcoded oligonucleotides released. These oligonucleotides have complementarity to sequence on ATAC fragments downstream of the sample index, which can lead to fragment swapping within multinucleated droplets. To circumvent this, we employed two complementary strategies: first, we added a blocking oligonucleotide in excess to the droplet barcoding step with a 3’-inverted dT-end. These blocking oligonucleotides compete with the carry-over Tn5-barcodes for binding of ATAC fragments. Their 3’-inverted dT-modification prevents further amplification of these fragments, preserving the balance of ATAC and RNA fragments and the correct sample barcodes in the library. Second, we decreased the number of thermal amplification cycles during droplet barcoding from twelve to four. By reducing the number of linear amplification cycles, the probability of a carryover Tn5-barcode binding a fragment and erroneously barcoding and exponentially amplifying it decreases. For Chip loading, the standard 10x workflow was followed. In brief, 70 µl of the sample, 50 µl of Barcoding Gel Beads, 40 µl Partitioning Oil were loaded into the respective channels. After the Chromium run, the droplet emulsion was transferred to 10x verified PCR tubes (Eppendorf, catalog no. EP0030124359) and the cell barcoding reaction was run in the thermocycler as follows: 5 min at 72 °C, 30 seconds at 98 °C, 12 cycles (98 °C 10 seconds, 59 °C 30 seconds and 72 °C 1 min). In the NIH-3T3/K562 species mix experiment, the droplet emulsion was split into two fractions to test 12 versus 4 cycles of linear amplification, and 20 % of each fraction was further processed to limit the sequencing requirements. Sample cleanup was performed following the standard 10x Genomics protocol with minor modifications: 125 µl Recovery Reagent was added to sample and inverted 10-15 times. After brief centrifugation, 125 µl of the pink oil phase was discarded and the aqueous phase was mixed with a dynabead cleanup mastermix (182 µl Cleanup Buffer, 13 µl Dynabeads MyOne SILANE and 5 µl Reducing Agent B (10x Genomics)) and incubated for 10 min at RT. Samples were placed on a magnetic separator and washed 2 times with 80 % ethanol before eluting them in 42 µl of elution buffer. Finally, samples were subjected to a 1.4x SPRI cleanup (Beckman Coulter, catalog no. B23318).

#### Pre- and final library amplification

Following GEM generation, cell barcoding and cleanup, samples containing a mixture of single-cell ATAC and RNA libraries were resuspended in the pre-amplification mastermix containing 1x NEBNext HF 2x PCR Master Mix (NEB, catalog no. M0541L) and a combination of three primers: partial-P5-PTO primer (common for both modalities), TruseqR2 (specific for the RNA modality) and an ATAC-spacer primer (specific for ATAC modality) each at a final concentration of 500 nM. Samples were incubated in the thermal cycle as follows: 98 °C 1 mi, 5 cycles of 98 °C for 20 sec, 63 °C for 20 sec and 72 °C for 20 sec), 72 °C for 1 min and then subjected to a 1.2x SPRI cleanup eluting in 42 µl EB buffer (Qiagen). Each sample was split into two fractions and each of them was further amplified with modality-specific primers: a partial-P5-PTO/P7-index-spacer primer pair for the ATAC libraries and a partial-P5-PTO/P7-index-TruseqR2 primer pair for the gene expression libraries. The PCR programme for both library types was as follows: 98 °C for 1 min, 5 cycles of 98 °C for 20 sec, 63 °C for 20 sec and 72 °C for 20 sec followed by a final extension step at 72 °C for 1 min. To estimate the number of endpoint PCR cycles required for each library, 1/20^th^ of the total library volume was used to perform a qPCR reaction using library-specific primers, and the number of cycles required to reach 50 % of saturation was calculated. Finalized ATAC libraries were subjected to a 1.2x SPRI cleanup and RNA libraries to 0.72x SPRI cleanup. The concentration of the libraries was measured using Qubit 1x dsDNA-HS (Thermo Scientific, catalog no. Q33230) and the library fragment distribution was assessed using Bioanalyzer High Sensitivity DNA kit (Agilent, catalog no. 5067-4626).

#### Sequencing

Libraries were sequenced on the NovaSeq6000 (Illumina) using a 100-cycle S2 Kit or NextSeq2000 (Illumina) using a 100-cycle P3 kit (ATAC: Read 1: 55 cycles, Index 1: 11 cycles, Index 2: 16 cycles, Read 2: 55 cycles, RNA: Read 1: 95 cycles, Index 1: 6 cycles, Index 2: 16 cycles, Read 2: 21 cycles).

### Data analysis procedures

#### ATAC sequencing data preprocessing

Base calls were converted to fastq format and demultiplexed by i7, allowing for one mismatch, using bcl-convert (v4.0.3). Demultiplexed reads were aligned to the hg38 genome and fragment file generation was performed with chromap (v0.2.3)^60^. For the species mixture experiment to evaluate collision rates, a combined hg38/mm10 reference was used and fragments aligning to either genome were counted. Using the R package ArchR (v1.02)^61^, generated fragment counts for each cell were computed in 500 base genome bins. The basic preprocessing pipeline for data processing until the ArchR object generation is available at: https://git.embl.de/grp-zaugg/SUMseq (available upon publication).

Cell barcode filtering was performed based on number of unique fragments, TSS enrichment score, and fraction of reads in promoter regions for each sample. Peak calling was performed for each sample separately. In brief, MACS2 peak caller is used to identify peaks for each group of cells, whereafter an iterative overlap peak merging procedure is done to derive a consensus peakset between all samples. BigWig files for trace plots were generated with ArchR::plotBrowserTrack normalizing for reads in TSS.

#### M1 and M2 signature score generation

To determine M1 and M2 signature genes, we obtained publicly available datasets and used DESeq2^62^ to identify genes that were differentially expressed in M1 and M2 macrophages compared to M0 (adjusted p-value < 0.05 and fold change > 0). The following datasets were utilized: GSE159112^63^, where THP1-derived macrophages were stimulated with LPS and IFN-γ or IL-4 and IL-13 for 24 hours to obtain M1 and M2 macrophages, respectively; and GSE55536^64^, involving iPSC-derived macrophages and human monocyte-derived macrophages stimulated with IFN-γ and LPS for M1 polarization, or IL-4 for M2 polarization. Genes identified as differentially expressed in both datasets were used as genesets for input to the ranking based scoring-approach AUCell^65^.

#### Transcription factor motif accessibility

Position weight matrices (PWMs) of TF-binding motifs were obtained from HOCOMOCO v12 (*in vivo* subcollection)^66^. Motif positions in the accessible chromatin regions were determined using the function ArchR::addMotifAnnotations. Z-score of bias-corrected (GC content and mean peak accessibility) per-cell motif accessibility was calculated with the function ArchR::addDeviationsMatrix, which leverages the chromVAR framework^27^.

#### RNA sequencing data preprocessing

Base calls were converted to fastq format using bcl2fastq (v2.20.0) or in the case of multiple pooled libraries base calls were converted to fastq format and demultiplexed by library index (i7) using bcl-convert (v4.0.3). The cell barcode (i5) was concatenated to the sample index and UMI (Read2), whereafter reads were demultiplexed by sample index using Je (v2.0.RC), allowing for two mismatches. Read alignment to Hg38 and gene expression matrix generation for each demultiplexed sample was conducted with STARsolo. For the species mixing experiment, a combined Hg38/mm10 reference was used and number of UMIs aligning to each genome was determined for each cell to estimate collision rates. For each sample, cell calling was either performed with EmptyDrops (v.1.16) or by determining inflection points on a rank vs UMI plot. Additionally, cells were filtered by mitochondrial and ribosomal read percentage. Feature by cell matrices were merged between samples, and finally features found in a minimum of 10-25 cells were retained.

#### Comparison to other technologies

We compare the performance of SUM-seq to dsci-ATAC-seq^11^GSM3507387), SHARE-seq^14^GSM4156590, RNA: GSM4156602 and GSM4156603), Paired-seq^15^ (ATAC: GSM3737488, RNA: GSM3737489), and scifi-RNA-seq^9^ (GSM5151362) using cell line data. We used NIH-3T3 data for SUM-seq, sequenced to approximately 30.000 reads/cell for each modality. For other methods, we use authors’ count matrices provided in the indicated repositories.

#### Dimensionality reduction and modality co-embedding

The peak matrix was normalized using TF-IDF, whereafter singular value decomposition was applied to derive a low-dimensional representation. The number of components considered was based on the proportion of variance explained (range 30-50), discarding the first component from downstream analysis due to high correlation with number of fragments per cell.

The gene expression matrix was normalized by proportional fitting (cell depth normalization to the mean cell depth), followed by logarithmic transformation (log(x + 0.5)), and another round of proportional fitting^67^. Next, a low dimensional representation of the normalized gene expression matrix was determined with PCA. The number of components considered was based on the proportion of variance explained (range 30-50).

Derived low-dimensional representations for each modality was used as input to learn cell modality weights, from which a WNN graph was constructed^68^. The WNN graph was further used to define a common UMAP visualization of the data modalities.

#### Multiomic factor analysis

To generate a common low-dimensional latent space for ATAC and RNA in the M1 and M2 polarizations, we utilized the multiomics integration platform MOFA^24^. First, ATAC data was collapsed from peaks to cis-regulatory topics with cisTopic (v0.3)^69^, determining a suitable number of topics based on the maximum on the second-derivative of likelihood curve and minimum on the model perplexity curve. All topics were used as input for MOFA. For RNA, top 4000 (default) most variable genes were used as input.

To determine polarization-associated latent factors, latent factors were correlated with biological metadata (state (M0/M1/M2), timepoint) as well as technical metadata (UMI/cell, fragments/cell). Factors with a high correlation against biological metadata, but not technical metadata were chosen as factors for downstream analysis.

#### Gene set enrichment analysis

Gene set enrichment analysis was performed for Reactome database genesets (v59). For every gene set G, significance is evaluated via a parametric *t*-test, where weights of the foreground set (features in set G) are contrasted against a background set (weights of features not in set G). *P* values were adjusted for multiple testing using the Benjamini–Hochberg procedure. Enrichments with false discovery rate 5 % were considered significant.

#### Motif enrichment analysis

Motif enrichment analysis was performed with R package monaLisa (v1.8) against HOCOMOCO v12 (*in vivo* subcollection) PWMs. Peaks within topics in the top and bottom 3 % quantile of feature weights for MOFA factors of interest were considered for analysis. TFs with FDR 0.1 % and log2 enrichment in the top or bottom 10 % quantiles were considered significant.

#### Gene regulatory network inference and transcription factor prioritization

We inferred an enhancer-based gene regulatory network (eGRN) using GRaNIE^36^ v1.5.3 with TFBS predictions based on HOCOMOCO v12^66^ database (*in vivo* subset) that were generated using PWMScan (see^70^ for methodological details how these were produced). The single-cell data from all time points and stimulations was first clustered using the smart local moving (SLM) algorithm with a resolution of 1, giving rise to 31 clusters that have at least 25 cells.

These clusters largely align with the stimulations and time points (**Extended Data Figure 5a**). We then calculated the mean RNA and ATAC values for each of these clusters as a pseudobulk, and input these clusters as ‘samples’ to create a eGRN following the GRaNIE single cell vignette (https://grp-zaugg.embl-community.io/GRaNIE/articles/GRaNIE_singleCell_eGRNs.html). We kept links if the TF-peak connection had an FDR < 0.2 and the peak-gene connection had an FDR < 0.1, the defaults in the GRaNIE package. We collapsed TF motif connections to the level of TFs to construct TF regulons.

#### Residual analysis

To identify discrepancy and concordance between TF motif accessibility and regulon activity scores, values were rank min-max normalized and the difference between the two was computed, and is referred to as the residual value. AUCell was used to infer single-cell activity scores for regulon geneset

#### GWAS integration

##### LDSC

To integrate GWAS with our eGRN, we performed stratified linkage disequilibrium score regression (S-LDSC^43^ v1.0.1). We added all open regions in all macrophage cells as background, on top of the default baseline model that includes genic regions, enhancer regions, and conserved regions. We then tested enriched heritability for all peaks that were part of the STAT1/STAT2/IRF9 eGRN, all peaks in the eGRN, and all peaks in M1 and M2 cells only. We included 8890 traits from UK biobank, FinnGen and several GWAS repositories as included in the 22.10 release of Open Targets Genetics^71^. SNP heritability (h2) was calculated with defaults from the LDSC codebase. We tested S-LDSC only if h2 was > 0.05, resulting in a list of 1230 traits. We corrected the enrichment p-values for the number of peaksets tested in each trait.

##### Gene modules and SNP overlap

For those traits that were enriched for heritability (adjusted p < 0.1), the SNPs associated to the particular trait (suggestive GWAS p-value < 5*10^-6) were intersected with the peaks in the STAT1/STAT2/IRF9, and full eGRNs (Supplementary Tables 7-8). We selected the genes connected (in the STAT1/STAT2/IRF9 eGRN) to the peaks that overlapped a SNP to obtain disease gene modules for CD, IBD and eosinophil counts.

## Code availability

A Snakemake preprocessing pipeline for SUM-seq data is available in a public Git repository (https://git.embl.de/grp-zaugg/SUMseq; available upon publication). All analysis source code will be available on a separate Git repository.

## Data availability

All data will be deposited to the NCBI GEO (GSE253165) or EGA databases. The provided links can be utilized to privately access the datasets before their public release. Data related to the species mixing experiment is available at GEO with accession number GSE253165 (https://www.ncbi.nlm.nih.gov/geo/query/acc.cgi?acc=GSE253165) and reviewer token upyfaouoxzgxvsx.

Processed data related to the iPSC-derived macrophage polarization experiment is available at: https://drive.google.com/drive/folders/1Y-

WM8fKMSaIk1iPzZ9jSDsBgf3lunWjG?usp=share_link (raw data will be deposited at EGA).

## Acknowledgements

We thank the EMBL Genomic Core facility for help with sequencing, Protein Purification Core facility for Tn5 and RNasin production, Michael Snyder for providing hiPSCs, Yakov Tsepilov and Xiangyu (Jack) Ge for providing access to the cleaned and harmonized database of GWAS summary statistics, and EMBL IT for access to the HPC cluster used for all analyses. We further thank members of the Zaugg and the Noh group for extensive discussions, Nadine Fernandez-Novel Marx for experimental support, and Josephine Brysting for providing cell lines.

A.C. was supported by an EIPOD4 Fellowship from the EC Horizon2020 MSCA (grant agreement number 847543). N.S. and J.B.Z. acknowledge funding from GSK through the EMBL-GSK collaboration framework (3000032294). N.H.S. and J.B.Z. acknowledge funding from the EMBL Infection Biology Transversal theme. K.D.P. was supported by SNSF (P2ZHP3_199669) and EMBO (ALTF538) Postdoctoral Fellowships. This work was supported by the Cariplo foundation grant, the GSK basic research fund, and the EMBL research fund (to K.M.N.). M.M. was supported by the Research council of Finland (grant number 347543), Sigrid Jusélius foundation, and Instrumentarium Science foundation. This project is co-funded by the European Union (ERC, EpiNicheAML,101044873) to J.B.Z.. Views and opinions expressed are however those of the author(s) only and do not necessarily reflect those of the European Union or the European Research Council. Neither the European Union nor the granting authority can be held responsible for them.

## Author Information

These authors contributed equally: Sara Lobato-Moreno, Umut Yildiz, and Annique Claringbould; position of S.L. and U.Y. was determined by coin flipping

## Contributions

M.M. conceived the project with input from U.Y., K.M.N. and J.B.Z. M.M., S.L. and U.Y. established and implemented the protocol and conducted experiments with the assistance of H.G.B., K.D.P. and V.C. M.M. and A.C. performed computational analyses. C.A. and E.P.V. constructed the pre-processing pipeline. N.H.S. assisted in data interpretation and manuscript writing. S.L., U.Y., A.C., M.M., J.B.Z. and K.M.N wrote the manuscript with input from all co-authors. M.M., J.B.Z. and K.M.N. supervised and directed the project. J.B.Z., K.M.N and M.M. secured financial support.

## Corresponding authors

Correspondence to Kyung-Min Noh, Mikael Marttinen or Judith B. Zaugg

## Ethics declarations

hiPSCs used for macrophage generation were derived from peripheral blood mononuclear cells with institutional review board approval (Stanford University, reference numbers 29904, 30064). The use of hiPSCs was approved by the EMBL Research Ethics Committee.

