## Extended Data Figures for "Scalable ultra-high-throughput single-cell chromatin and RNA sequencing reveals gene regulatory dynamics linking macrophage polarization to autoimmune disease"

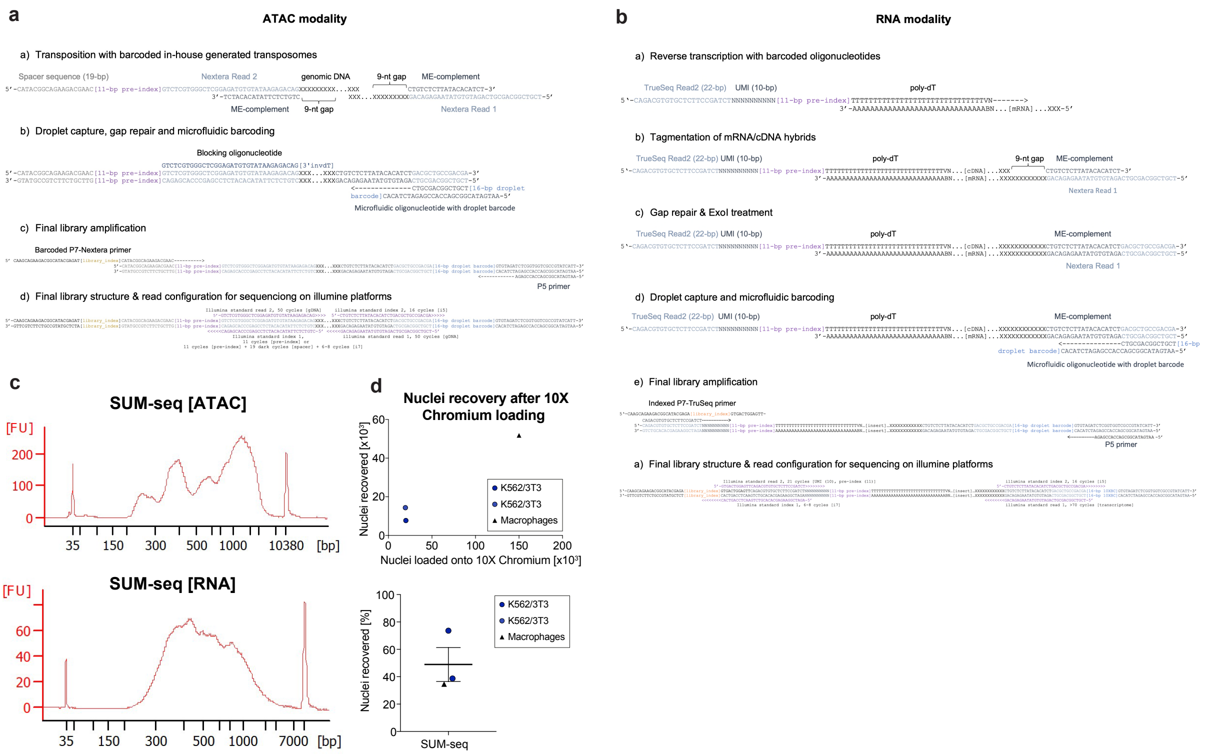

### Extended Data Figure 1. Detailed library structures of SUM-seq and performance metrics

**a**, Detailed outline with oligonucleotide sequences of the ATAC modality of SUM-seq for each step of the experimental workflow. **b**, Detailed schematic outline with oligonucleotide sequences of the RNA modality of SUM-seq. **c**, Bioanalyzer profiles of final SUM-seq snATAC- (upper panel) and snRNA-seq (lower panel) libraries. **d**, Raw recovery values (upper panel) and percentages (lower panel) of nuclei recovery before and after 10x Chromium loading.

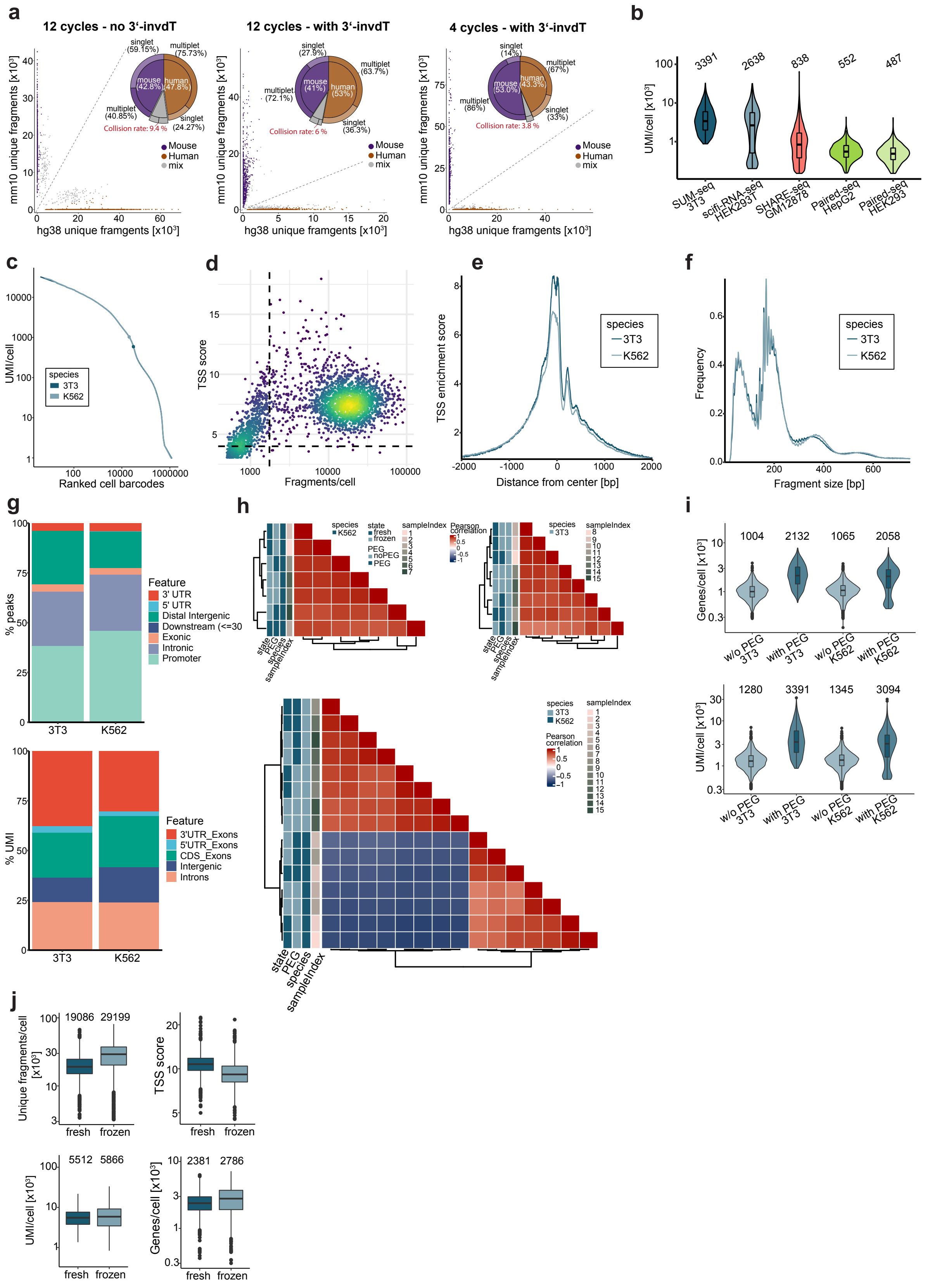

**Extended Data Figure 2. Key performance metrics of SUM-seq and optimization of experimental parameters**

**a**, Species-mixing experiments of human K562 and mouse NIH-3T3 cells with SUM-seq and the impact of the blocking oligonucleotide and reduction of thermal amplification cycles on collision rates (highlighted in red). The y-axes indicate the number of reads derived from each cell mapped to the mouse genome (mm10), while the x-axes show the number of reads for each cell mapped to the human genome (hg38). The pie chart shows the fraction of human and mouse cells as well as the frequency of multiplets and singlets. **b**, UMIs detected per cell for SUM-seq, scifi-RNA-seq, SHARE-seq and Paired-seq in the indicated cell lines (datasets used for the comparison are the same as in **Fig. 1e** (lower panel). **c**, Knee plot showing the number of UMIs (y-axis) per cell ranked according to frequency (x-axis) of the species-mixing SUM-seq experiment. The inflection points are indicated as points along the curve. **d**, Scatterplot displaying the TSS score (y-axis) and the number of fragments (x-axis) per cell for the ATAC modality of the NIH-3T3 cells in the species-mixing SUM-seq experiment. Thresholds for cell calling are indicated with dashed lines. **e**, TSS-enrichment plots of snATAC-seq data for human K562 (lighter blue) and mouse NIH-3T3 (darker blue) cell lines. The enrichment of ATAC-reads (y-axis) is displayed around the TSS (±2 kb). **f**, Size distribution of ATAC fragments in the indicated cell lines in base pairs (x-axis). **g**, Genomic annotation of mapped ATAC fragments (upper panel) and transcripts (lower panel) in K562 and NIH-3T3 cells. **h**, Correlation heatmaps of sample indices for the ATAC- (upper panels) and RNA-modalities (lower panel) of the species-mixing SUM-seq experiment. Rows are annotated according to their state (fresh or frozen), cell type (K562, NIH-3T3), inclusion of PEG for the reverse transcription reaction and the utilized sample index. **i**, Violin plots displaying the effect of PEG on the number of genes (upper panel) and UMIs (lower panel) detected in the RNA modality of SUM-seq in NIH-3T3 and K562 cells. **j**, Impact of cryopreservation on the data complexity of SUM-seq. The upper panel displays key quality metrics comparison for the ATAC-modality (fragments per cell and TSS enrichment score), and the lower panel for the RNA modality (genes and UMIs detected per cell)

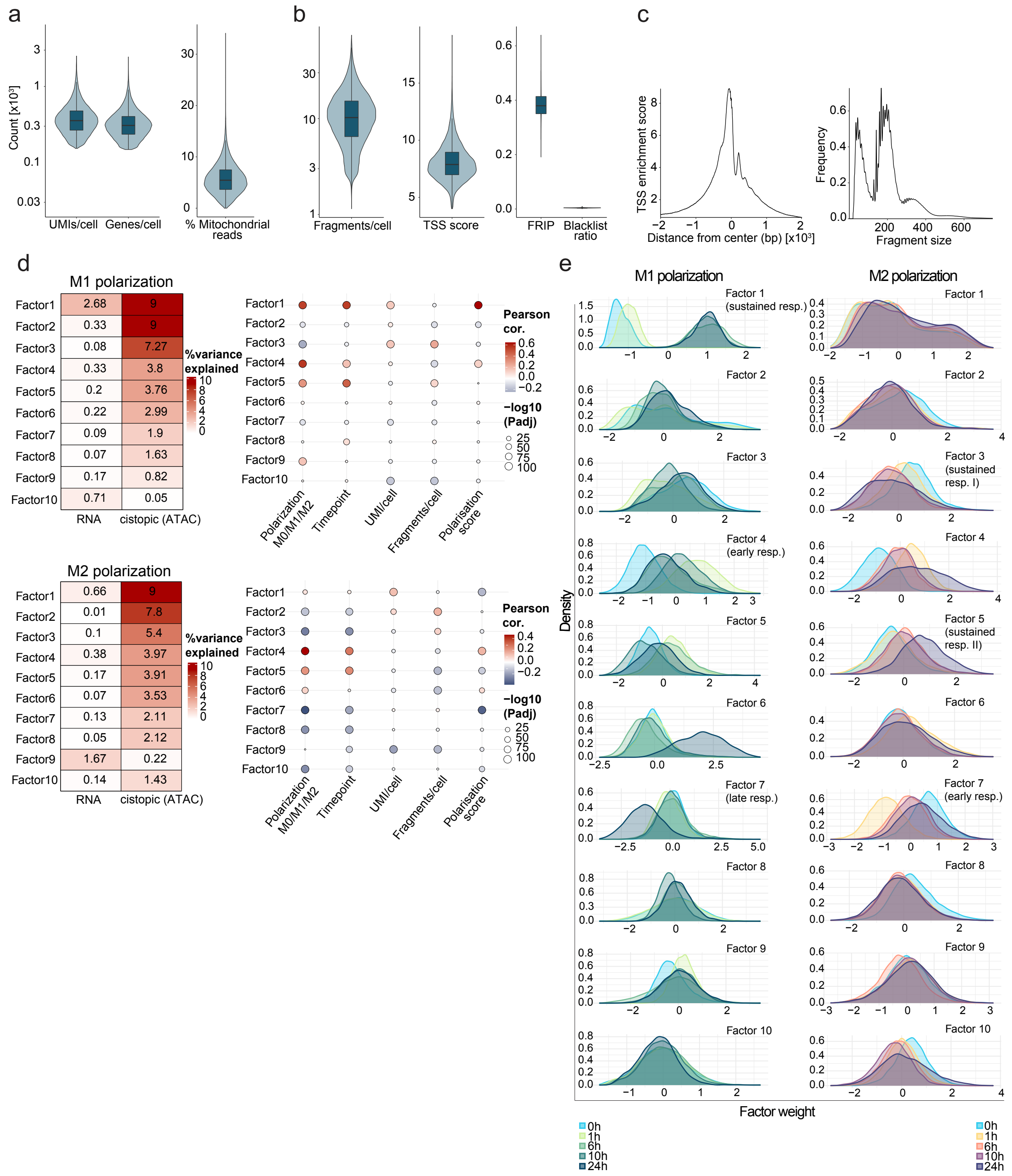

**Extended Data Figure 3. Quality control of M1/M2 polarization SUM-seq experiment and relations of latent factors with metadata**

**a,** UMIs, genes and percentage of mitochondrial reads per cell for the RNA modality, and **b,** unique fragments, TSS enrichment score, and fraction of reads in peaks (FRIP) per cell for the ATAC modality of the M1/M2 polarization SUM-seq experiment. **c,** The enrichment of ATAC-reads (y-axis) displayed around the TSS (±2 kb) (left). Size distribution of ATAC fragments in the indicated cell lines in base pairs (x-axis) (right). **d,** The percentage of variance explained by MOFA-inferred latent factors for each modality (left) for M0/M1 cells (top) and M0/M2 cells (bottom). The correlation between latent factors and metadata (right) for M0/M1 cells (top) and M0/M2 cells (bottom). **e,** Distributions of time point cells across MOFA factor weights associated with M1 polarization (left) and M2 polarization (right).

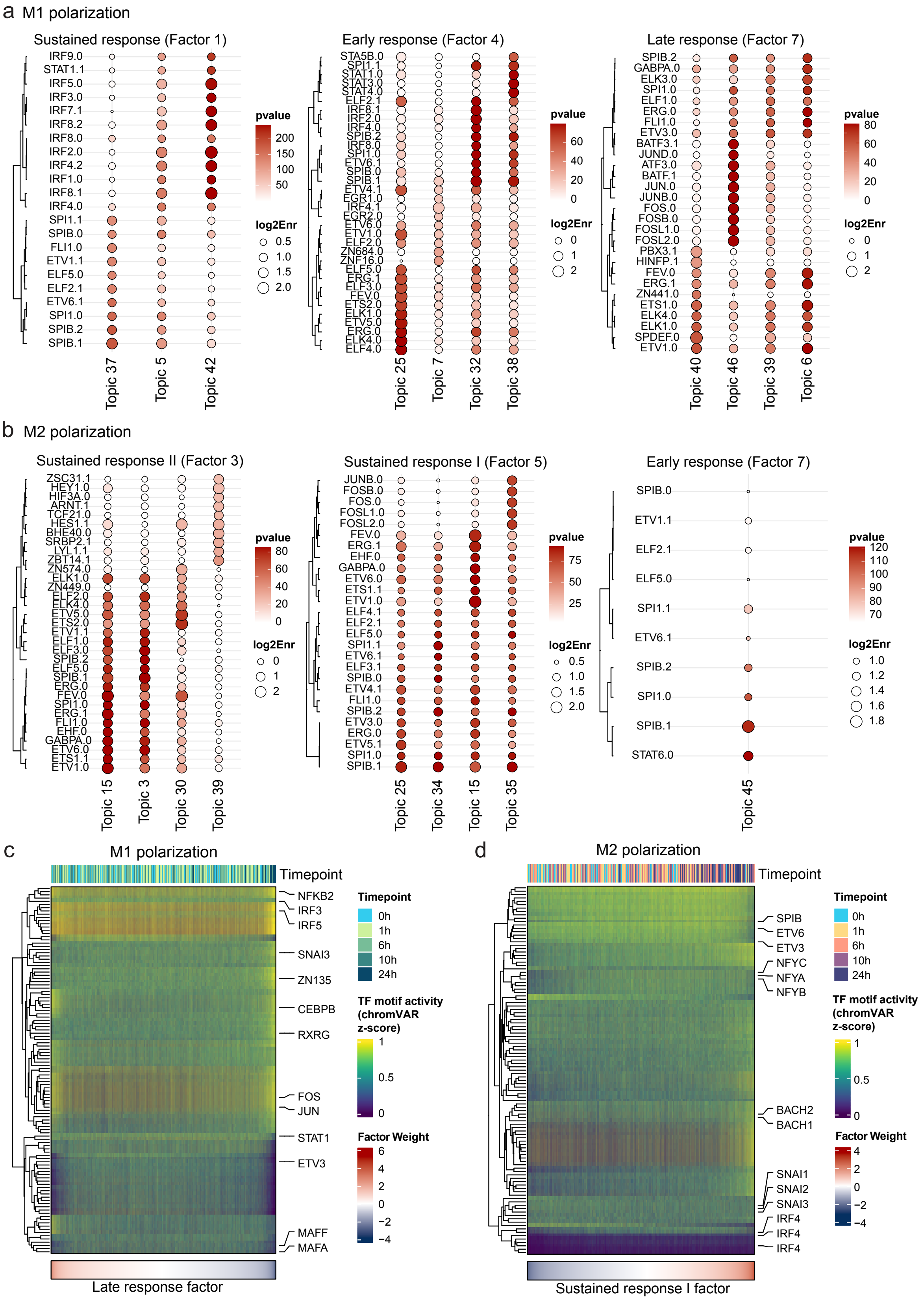

**Extended Data Figure 4. Motif enrichment for peaksets associated with M1/M2 latent factors and TF motif activity for M1 late response- and M2 sustained response II TFs**

**a,** Top 10 TF motifs enriched per topic peaksets associated with M1 sustained response (left), M1 immediate response (center) and M1 late response (right) factors. **b,** Top 10 TF motifs enriched per topic peaksets associated with M2 sustained response I (left), M2 sustained response II (center) and M2 immediate response (right) factors. **c,** Motif activity for TFs associated with M1 late response (**Methods**) across all M0 and M1 cells sorted by M1 late response factor weights **d,** Motif activity for TFs associated with M2 sustained response II across all M0 and M2 cells sorted by M2 sustained response II factor weights

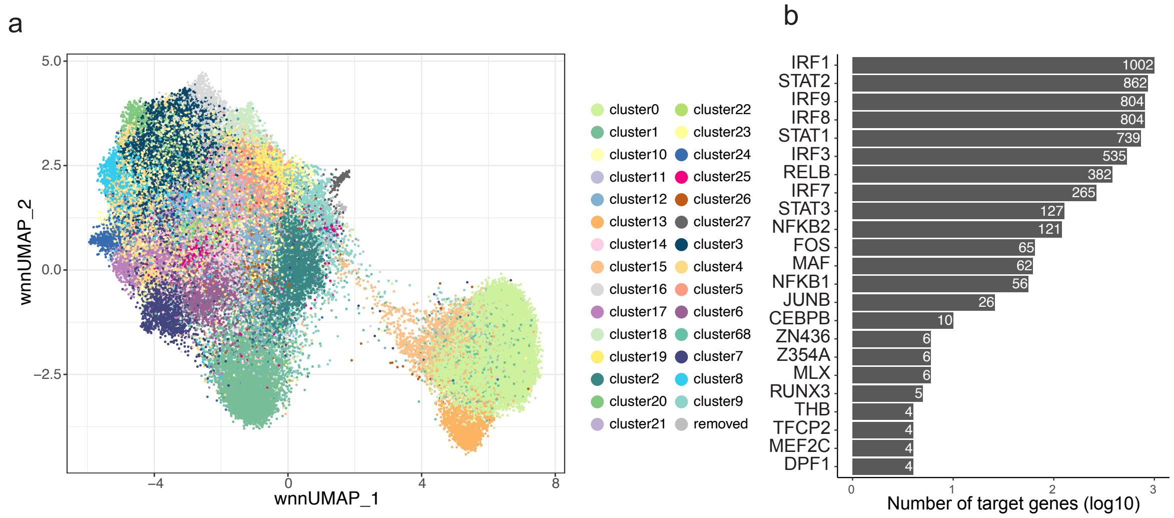

**Extended Data Figure 5. Gene regulatory network**

**a**, UMAP of the cell clusters used for pseudobulking and subsequently inferring a eGRN. Only clusters with more than 25 cells were included to make the eGRN. **b**, Degree plot Bar plot plot of number of target genes linked to each transcription factor (TF) in the eGRN. Only showing TFs with at least 4 connected genes.

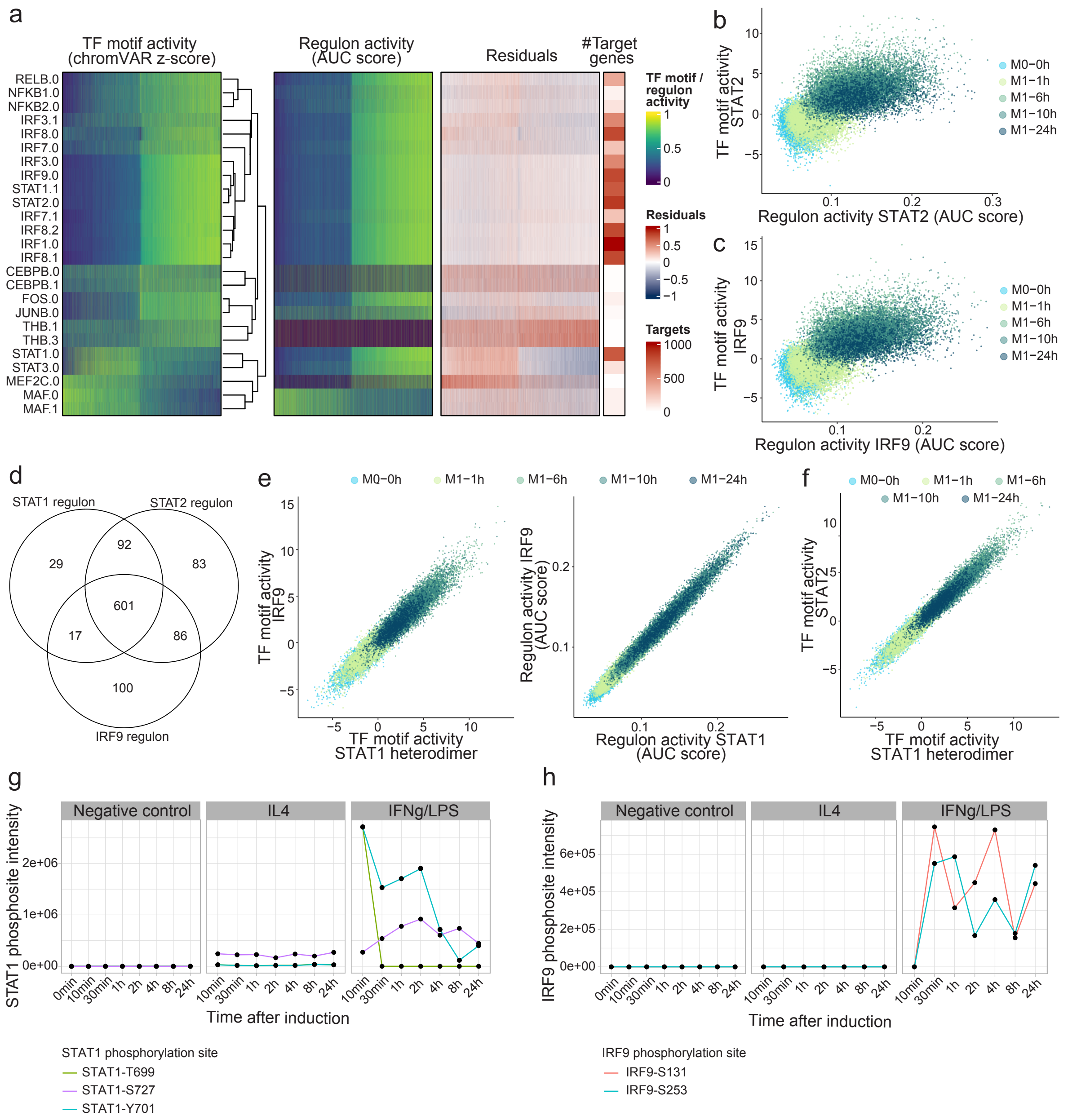

**Extended Data Figure 6. Dynamics of STAT1, STAT2, and IRF9 motif and regulon activity in M1 polarization**

**a,** Heatmaps showing TF motif activity (ChromVAR z-score), regulon activity (AUC score), and the similarity between the two, for M1 polarization-associated TFs found as TFs in the eGRN, across M0 and M1 cells sorted by the M1 sustained factor. **b,** Scatter plots of the correlation between the STAT2 regulon activity (AUC score) and TF motif activity (chromVAR Z-score) and **c,** IRF9 regulon activity and motif activity. Points represent cells and are coloured by their experimental time point. **d,** Venn diagram of STAT1, STAT2 and IRF9 regulon genes. **e,** Scatter plots of the correlation between STAT1 heterodimer and IRF9 motif activities (top) and STAT1 and IRF9 regulon activities (bottom). Points represent cells and are coloured by their experimental time point. **f,** Scatter plots of the correlation between STAT1 heterodimer and STAT2 motif activities (top) and STAT1 and STAT2 regulon activities (bottom). Points represent cells and are coloured by their experimental time point. **g,** STAT1 T699 and Y701 phosphopeptide intensities upon IFN-y/LPS stimulation of THP1 derived macrophages. **h,**

IRF9 S131 and S253 phosphopeptide intensities upon IFN-y/LPS stimulation of THP1 derived macrophages.

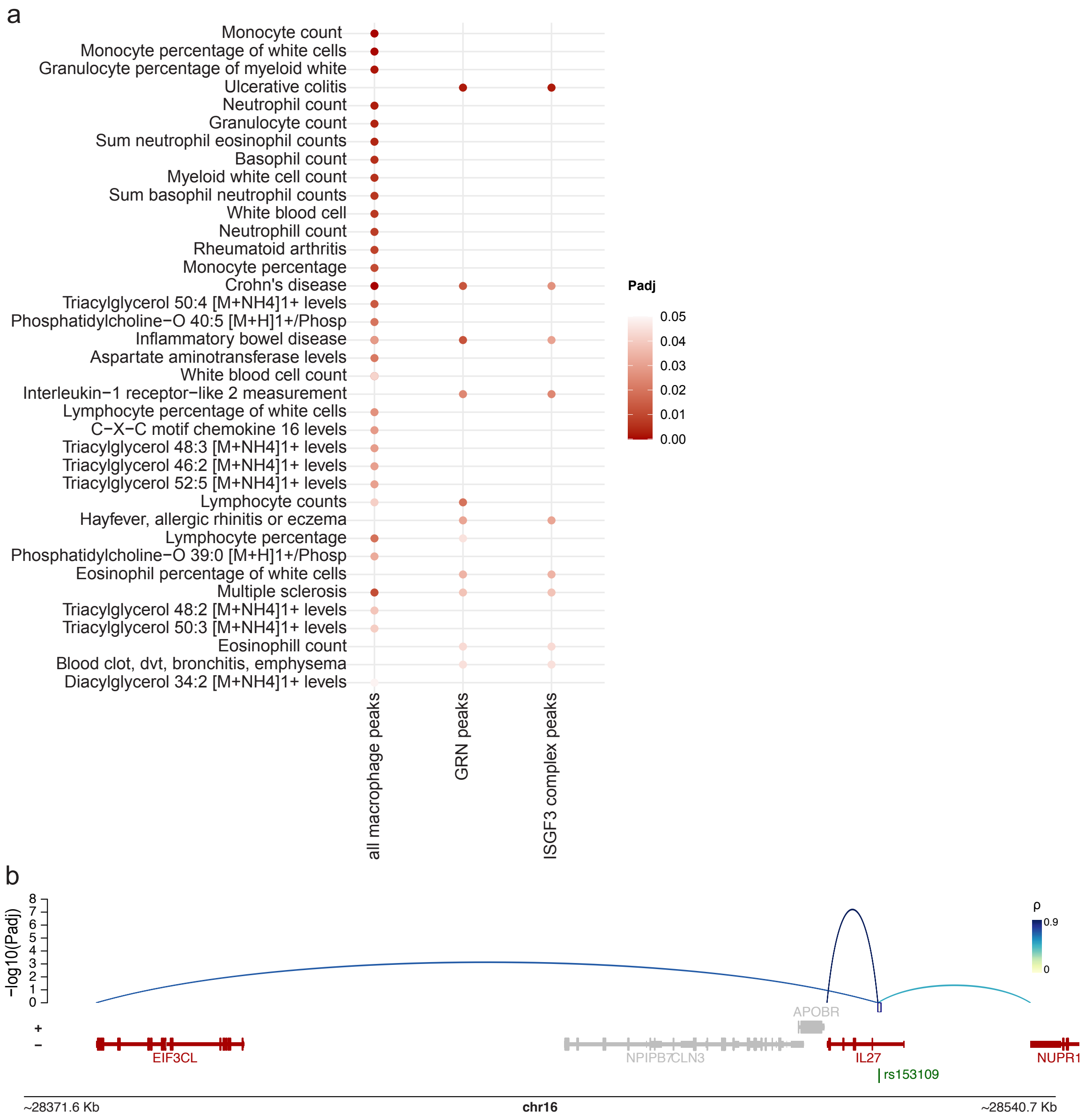

**Extended Data Figure 7. Extended LDSC results**

**a,** LDSC for all eGRN peaks. **b, e**GRN peak-gene interactions for the peak upstream of CHI3L1 gene intersecting with SNP rs872129 (top) and M0 and M1 cell aggregate ATAC-seq tracks split by time point (bottom).

### Supplementary Tables 1-7:

**Supplementary Table 1:** SUM-seq oligonucleotide sequences
**Supplementary Table 2:** M1 and M2 marker genes used for determining M1 and M2 scores
Column names:
Polarisation = *M1 or M2 polarisation*
Gene = *marker gene name*
**Supplementary Table 3:** Gene set enrichment analysis (GSEA) for M1 and M2 early, sustained and late response factors
Each tab contains the GSEA for M1 or M2. Column names:
Reactome pathway = pathways of the Reactome database
Factor = adjusted p-values for GSEA of gene sets associated with selected MOFA factors (associated with early, sustained and late M1 and M2 responses)
**Supplementary Table 4:** Top variable transcription factor (TF) motifs identified by ChromVAR and and motif enrichment in M1 and M2 MOFA (Multi-Omics Factor Analysis) factors
Tabs 1 and 2 contain the top 10% most variable TF motifs in M1/M2 polarization and filtered by those enriched in early, sustained and late response factors
Tabs 3 and 4 -log10 adjusted p values for motif enrichment analysis on cistopics associated with M1-polarisation relevant latent factors. Columns are cistopics and rows are motifs.
Tabs 5 and 6 log 2 enrichment for motif enrichment analysis on cistopics associated with M1-polarisation relevant latent factors. Columns are cistopics and rows are motifs.
**Supplementary Table 5:** Gene regulatory network (GRN) connections, filtered for TF-peak FDR =< 0.2 and peak-gene FDR =< 0.1
Column names:
TF.ID, TF.name, TF.ENSEMBL = *Transcription factor motif, name and ENSEMBL ID*
peak.ID = *Location of the peak (chromosome:start-end)*
peak.mean, peak.median = *Mean and median accessibility of peak*
peak.CV = *Coefficient of variation of accessibility of peak*
peak.annotation = *Type of peak (e.g. intronic, promoter, intergenic)*
peak.nearestGene.chr, peak.nearestGene.start, peak.nearestGene.end = *Location of the nearest gene from this peak (chromosome, start, end)*
peak.nearestGene.length, peak.nearestGene.strand, peak.nearestGene.distanceToTSS = *Length, strand and distance to transcription start site of nearest gene from this peak*
peak.nearestGene.name, peak.nearestGene.ENSEMBL, peak.nearestGene.symbol = *full name, ENSEMBL ID and name of nearest gene from this peak*
peak.GC.perc = *Percentage of GC content in peak*
TF_peak.r, TF_peak.r_bin = *Correlation coefficient and bin between TF and peak*
TF_peak.fdr, TF_peak.fdr_direction = *False discovery rate and direction of effect of correlation between TF and peak*
TF_peak.connectionType = *Method used to calculate TF activity*
peak_gene.distance = *Distance between peak and target gene*
peak_gene.p_raw, peak_gene.p_adj = *Raw and adjusted p-value for correlation between peak and target gene*
gene.ENSEMBL, gene.name = *ENSEMBL ID and name of target gene*
gene.type = *Type of target gene (e.g. protein coding, lncRNA)*
gene.mean, gene.median = *Mean and median expression of target gene*
gene.CV = *Coefficient of variation of expression of target gene*
gene.chr, gene.start, gene.end = *Location of target gene (chromosome, start, end)*
gene.strand = *Strand of target gene*
TF_gene.r, TF_gene.p_raw = *Raw p-value and correlation coefficient between TF and target gene*
**Supplementary Table 6:** STAT1 regulons from the GRN

Columns: STAT1 motif (first column), target gene name (second column)

**Supplementary Table 7:** Disease SNP overlap with all GRN regulons

Column names:

SNP = *Single nucleotide polymorphism rs ID*

peak = *GRN peak that overlaps with the SNP*

SNP.Z = *GWAS Z-score for the SNP corresponding to A1 allele*

SNP.A1, SNP.A2 = *Reference (A1) and alternative (A2) SNP alleles*

TF.name, TF.ENSEMBL = *Name and ENSEMBL ID for TF linked to peak*

gene.name, gene.ENSEMBL = *Name and ENSEMBL ID for gene linked to peak*

gene.chr, gene.start, gene.end = *Location of target gene (chromosome, start, end)*

peak_gene.p_adj = *Adjusted p-value for correlation between peak and target gene*

peak_gene.r = *Correlation coefficient between peak and gene*

disease = *GWAS trait where the SNP was identified (format: disease_studyID)*

**Supplementary Table 8:** Disease SNP overlap with STAT1/STAT2/IRF9 regulon

Column names as in Supplementary Table 7

**Supplementary Table 9:** Expression, protein, splicing quantitative trait loci (e-, p-, sQTLs) for rs4810485, collated by Open Targets Genetics

The tabs show results for eQTLs, pQTLs and sQTLs.

Columns indicate the dataset, rows the genes, values the significance as displayed on Open Targets website.
